# Minimally invasive monitoring of clonal evolution through integrated single cell and ctDNA analysis

**DOI:** 10.64898/2026.07.29.741620

**Authors:** Farhia Kabeer, Matteo Lepur, Branden J. Lynch, Emilia Hurtado, Elena Zaikova, Janine Senz, Vinci Au, Caroline Baril, Ding Ma, Sheila Nicholson, LUMES Consortium, Gavin Ha, Jessica N. McAlpine, Samuel Aparicio, David G. Huntsman, Alexandre Bouchard-Côté, Yvette Drew, Andrew J. L. Roth

## Abstract

Circulating cell-free DNA (cfDNA) offers a minimally invasive lens into tumor evolution. However, the accurate quantification of clonal composition from cfDNA remains challenging. Existing methods for clone tracking using liquid biopsies are constrained by issues such as their reliance on bulk tissue references, incomplete representations of clonal architecture or limited breadth of mutation panels. To address these limitations, we developed **cfClone**, a Bayesian framework that integrates single-cell whole-genome sequencing (scWGS) derived clonal structures with cfDNA whole-genome sequencing (cfDNA-WGS) data to enable high-resolution tissue-informed clonal tracking. By jointly modeling local copy-number alterations and haplotype-specific signals via Bayesian model selection and Markov chain Monte Carlo (MCMC) sampling, the algorithm yields uncertainty-aware estimates of clonal prevalence and tumour fraction (TF). Applied to longitudinal clinical cohorts, cfClone can be used to reconstruct evolutionary trajectories and uncovers clonal selection driving therapeutic resistance, including the *de novo* detection of emergent clonal populations. We demonstrate that cfClone achieves accurate TF estimates and circulating tumor DNA (ctDNA) detection in malignancies with varying degrees of copy-number variant (CNV) burden using semi-synthetic data. We then compare to the state of the art scWGS informed panel based approach, and demonstrate cfClone provides comparable accuracy while allowing for the tracking of more clones and detection of novel clones. Finally, we show how cfClone can be used to quantitatively track clonal dynamics in response to treatment in high grade serous ovarian cancer.

Github link: https://github.com/Roth-Lab/cfclone

## Main

Cancer is an evolutionary process, whereby clonal cell populations (clones) have varying degrees of fitness, which for example, can lead to selective resistance to treatments that change over time^1^. The ability to quantify the differential response to treatment between clones from the same patient, provides a potential opportunity to understand features associated with resistance, and in the clinical setting personalise treatment strategies. Tissue-based biopsies have been used to provide insight into cancer dynamics^2^, however obtaining multi-site and/or serial samples is often too invasive to be practical. Liquid biopsies are increasingly being considered as a viable method to obtain serial patient samples as they are minimally invasive, concordant to tissue biopsies, and show promise for early cancer detection^3–5^.

Cell-free DNA (cfDNA) derived from a blood-based liquid biopsy is the collection of the DNA fragments shed into the blood stream by cells undergoing apoptosis and necrosis^6^. These DNA fragments circulate throughout the blood stream from multiple sites in the body and different cells, providing a detailed overview of the patient’s cancer genome. A challenge when analyzing cfDNA, is that it is often dominated by DNA fragments originating from non-maligant cells, predominantly white blood cells, leaving a sparse signal from circulating tumour DNA (ctDNA)^7,8^. In principle, cfDNA serves as a heavily diluted tissue sample, albeit with specific biases that arise from differential rates of DNA degradation in blood due to various factors such as nucleosome occupancy, as well as the potential contribution of DNA from multiple tumor sites^17^.

Cancer alters the genome through a variety of mechanisms, including somatic copy number alterations (SCNAs), a process that copies and/or deletes large segments of the genome. As SCNAs are mainly found in tumour cells they provide a method to differentiate tumour derived DNA from normal DNA fragments enabling methods to track cancer progression and evolution^9^. Tumour cell populations can be further parsed into detailed subpopulations called clones based on their SCNAs profiles. This provides a better understanding of the tumour population itself, as it reveals tumour clones that drive treatment resistance and relapse. Existing methods for clonal analysis of cfDNA often leverage SCNAs and can be divided into two groups: those which incorporate prior information on clonal population structure, and those which do not. In the latter case, tools like ichorCNA^4^ yield estimates of tumor fraction, i.e., the fraction of cfDNA which is of tumor origin. Similarly, liquidCNA^10^ provides an estimate of tumor fraction, as well as estimating the prevalence of the dominant subclone in time-series cfDNA samples. Neither yield robust estimates of clonal population structure. Studies using bulk sequencing of tissue coupled with clonal deconvolution methods to resolve clones and define panels for cfDNA analysis have demonstrated the feasibility of tumour informed clonal tracking^11^. However, the use of bulk sequencing instead of single cell sequencing limits the fidelity and resolution of initial clone genotyping. We are aware of only one method (CloneSeq-SV) that uses prior information about clonal architecture derived from single cell sequencing to track clonal evolution in cfDNA samples. However, in addition to single cell whole genome sequencing (scWGS), CloneSeq-SV requires specialised error-corrected duplex sequencing and manually curated clone specific single nucleotide and structural variants (SNVs and SVs), which are used to generate patient-specific sequencing panels for cfDNA monitoring^12^. While powerful, the complexity and cost of implementing a patient-specific panel limits its general applicability and provides no way to detect clones that were not identified by scWGS. Furthermore, the use of clone specific SNVs and SVs may only be able to track a subset of clones detectable by SCNA analysis of scWGS.

To address these challenges, we developed a Bayesian method, cfClone, that jointly models copy number variation and B-allele frequency from cfDNA whole genome sequencing (cfDNA-WGS) using clone specific copy number profiles learned from scWGS, thereby improving sensitivity through integration of complementary signals and clone level resolution of tumour cell populations. cfClone employs Bayesian model comparison to provide a principled statistical framework for ctDNA detection. Applied to longitudinal samples, cfClone provides insight into tumour evolution and identifies therapy resistant clones. In addition, cfClone provides diagnostics to identify the presence of clones in cfDNA not identified by scWGS analysis.

## Results

The study design and cfClone workflow are summarized in Fig. 1a, b, including longitudinal clinical sampling (Fig. 1a) and integration of scWGS-derived clonal copy number profiles with read-depth ratio (RDR) and B-allele frequency (BAF) data obtained from cfDNA-WGS of plasma (Fig. 1b). cfClone uses scWGS of tumour samples to define haplotype specific copy number variants (HSCNVs) for each clone (Fig. 1c, d). Computational deconvolution of the combined RDR and BAF modalities obtained from cfDNA-WGS is performed with a Bayesian model, where the clone specific HSCNV profiles are used as prior information for the deconvolution. This yields estimates of clonal prevalence and tumor fraction (TF) in the cfDNA. To perform posterior inference, cfClone uses Markov chain Monte Carlo (MCMC)^13^ sampling. cfClone outputs posterior densities for model parameters such as clonal prevalence and tumour fraction allowing for estimation of uncertainty. A detailed description of the cfClone model, inference strategy, posterior summarisation and ctDNA detection are provided in supplementary note 1 (see also, Supplementary Fig. 1-2).

**Fig. 1.**
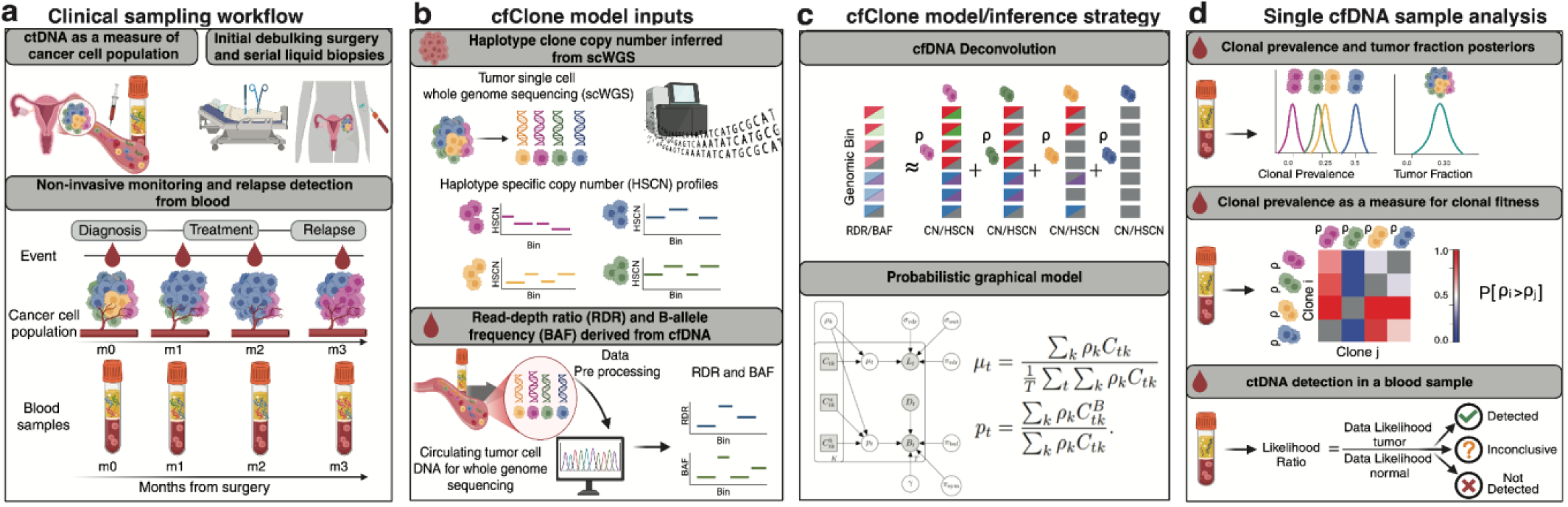
Methodological framework for minimally invasive tracking of tumor clonal dynamics using cfClone. Overview of the computational and clinical workflow **a**, Clinical sampling workflow tracking cancer cell populations and serial blood samples from initial debulking surgery through treatment and relapse **b**, cfClone data protocol integrating single-cell whole-genome sequencing (scWGS) clone copy number profiles with copy number variants (CNV) and B-allele frequencies (BAF) derived from circulating tumor DNA (ctDNA) **c**, cfClone model and inference strategy using a probabilistic graphical model for statistical cfDNA deconvolution **d**, Single cfDNA sample analysis outputs, detailing clonal prevalence, fitness evaluation, and baseline ctDNA detection metrics. **Abbreviations:** RDR, read-depth ratio; CN, copy number; RD, read depth; VAF, variant allele frequency; m, month; P, clone prevalence.

### Semi-synthetic mixtures of cell populations

We first evaluated the performance of cfClone using in-silico mixtures of scWGS data. Our simulation framework takes as input scWGS data, single cell clone assignments, tumour fraction, clone prevalences, sequencing coverage, and outputs binned total and haplotype read counts for a simulated bulk WGS sample (Supplementary note 2.1-2.2). We validated the framework by comparison to true cfDNA samples, ensuring that the simulated data recapitulated the observed cfDNA data when provided with the same tumour content and clonal prevalence values. Across 100 simulated data replicates, the 99% quantile interval of the simulated data contained 70% of the true cfDNA data in the RDR modality, and 99% of the true cfDNA data in the BAF modality (Supplementary Fig. 3). Using our simulation framework we then evaluated cfClone’s ability to infer tumour fraction, clonal prevalence, and detect ctDNA across a range of sequencing coverages, tumour fractions, and clonal prevalence values.

First we assessed the accuracy of cfClone’s tumour fraction estimation, and compared it with ichorCNA^4^, a widely used tumour naive CNV based method. We simulated samples across varying tumour fractions, levels of coverage. Simulations were performed using three cancer types with low (follicular lymphoma, FL), intermediate (diffuse large B cell lymphoma, DLBCL) and high (high grade serous ovarian cancer, HGSOC) levels of aneuploidy. cfClone and ichorCNA show comparable results when analysing FL data (30X coverage; cfClone, Pearson’s r = 1.00, p < 0.01, mean absolute error (MAE) = 0.243%; ichorCNA, Pearson’s r = 1.0, p < 0.01, MAE = 0.694%; Fig. 2a; Supplementary Fig. 4-5). cfClone outperformed ichorCNA when analysing data from DLBCL (30X coverage; cfClone, Pearson’s r = 1.00, p < 0.01, MAE = 0.195%; ichorCNA, Pearson’s r = 1.0, p < 0.01, MAE = 0.944%; Fig. 2a; Supplementary Fig. 6-7) and HGSOC (30X coverage; cfClone, Pearson’s r = 1.0, p < 0.01, MAE = 0.100%; ichorCNA, Pearson’s r = 1.0, p < 0.01, MAE = 1.57%; Fig. 2b; Supplementary Fig. 8-9)..

**Fig. 2.**
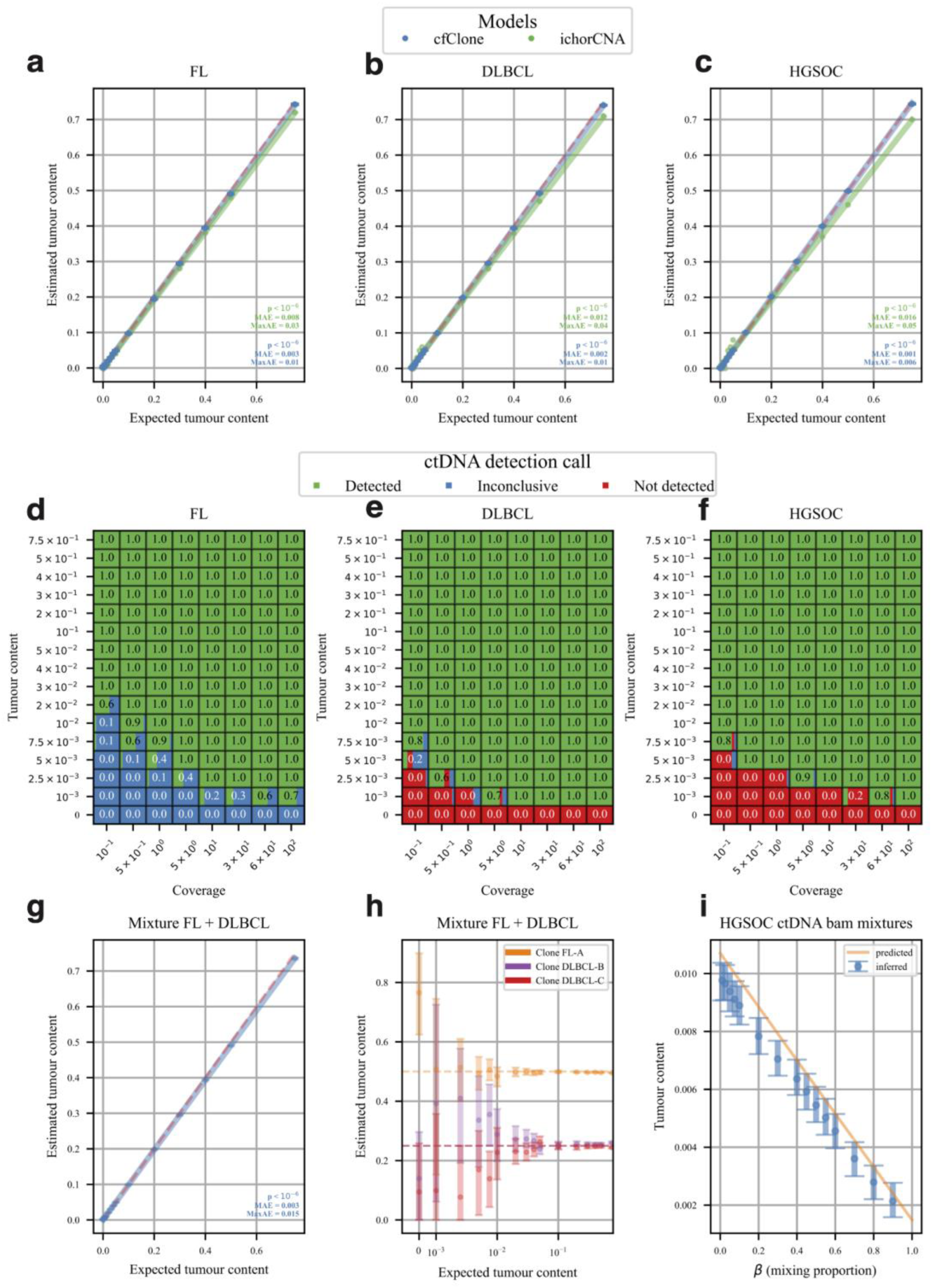
Synthetic benchmarking and in-silico validation. **a-c,** Benchmarking tumour content estimation of cfClone against ichorCNA with simulated cfDNA samples generated from the infinite pool model using scWGS of HGSOC, Diffuse large B-cell lymphoma (DLBCL) and Follicular lymphoma (FL) cancers. Ordinary Least Squares (OLS) regression was performed on the mean value cfClone’s posterior. OLS regression was performed on the most accurate solutions produced from ichorCNA’s multiple restarts. **d-e,** Proportion of ctDNA detections on cfDNA samples simulated using the infinite pool model with scWGS of HGSOC, FL, and DLBCL cancers. **g-h,** Tumour fraction and clone prevalence estimation of cfDNA samples generated by mixing scWGS of one FL clone and two DLBCL cancer clones at fixed proportions (0.5, 0.25, and 0.25, respectively); target coverage shown is 100X. **i,** Tumour content estimation by cfClone when mixing patient derived cfDNA samples with tumour content predicted by mathematical modeling (See Supplemental Note 3.3.2) .

Next we assessed cfClone’s sensitivity and specificity for binary detection of ctDNA across varying coverage, tumour fraction and levels of aneuploidy. Results for the complete grid of coverage and tumour fractions are summarised in Fig. 2d-f and Supplementary Figs. 10-12. For example, in the ultra low coverage (0.1x) and moderate tumour fraction (2%) simulations, cfClone detected ctDNA in 60% FL samples and 100% of DLBCL and HGSOC samples. In the high coverage (100X) coverage and low tumour fraction (0.1%) samples, cfClone detected ctDNA in 70% of FL samples and 100% of DLBCL and HGSOC samples.

Finally, we assessed cfClone’s ability to infer clonal prevalence by mixing clones from the FL and DLBCL tumours. We note these tumours are patient matched, having been obtained from an FL patient who underwent histological transformation to DLBCL^14^. We created in-silico mixtures by mixing 1 clone from the FL tumour with 2 clones from DLBCL tumour. The MAE of TF estimates was 0.300% at 100X target coverage (Fig 2g; Supplementary. Fig. 13). cfClone accurately deconvolved the FL/DLBCL mixtures (Fig. 2h; Supplementary. Fig. 13). In the low coverage setting (0.1X), across mixtures of tumor fractions ranging from 0.1% to 75%, cfClone estimated clonal prevalence for clone FL-A with an MAE of 6.06%, clone DLBCL-B with an MAE of 9.00%, and clone DLBCL-C with an MAE of 7.57%; increasing coverage to 100X reduced the MAE of clonal prevalence to 0.496%, 4.25%, and 4.15% respectively. The large decrease in MAE for clone A with increasing coverage is consistent with its distinct HSCNV profile (Supplementary Fig. 14). Conversely, the comparably small decrease in MAE for clones B and C is likely due to their similar HSCNV profiles (Supplementary Fig. 14), which leads to uncertainty about their prevalence due to model identifiability.

### Semi-synthetic mixtures of patient derived cfDNA samples

We next evaluated cfClone on a secondary simulation framework that accounts for cfDNA specific coverage biases arising from various factors, including differential rates of DNA degradation in plasma, nucleosome positioning, and other factors which may alter cfDNA abundance in a region or sequence specific fashion ^15^. To do so we generated data by mixing patient matched cfDNA or tissue samples in-silico at fixed proportions for a given target coverage (Supplementary note 2.3). We assessed two cfDNA sample combinations, in which a sample estimated to contain near zero tumor fraction (VOA12433P; TF = 0.149%) was mixed in proportions ranging from 0-0.9 with a sample containing low (VOA10703P; TF = 1.07%) or high (VOA8262P1; TF = 30.3%) estimated tumor fraction. We then compared the tumor fraction estimated by cfClone on the resulting mixtures to the expected theoretical tumor fraction predicted by our mathematical modelling. Similarly, we generated two mixtures of bulk tissue samples, derived from mixing one sample containing a low estimated tumor fraction (A78136; TF = 2.27%), with a low (VOA7648C; TF = 0.352%) or high (VOA7648A; TF = 79.8%) estimated tumor fraction sample, to further verify that our estimates are not confounded by cfDNA specific biases.

For a target coverage of 30X, results of the in-silico mixtures of cfDNA were comparable in accuracy to the mixtures derived from scWGS above, with a MAE of 0.0658% when mixing the near zero tumor fraction sample with the low tumor content sample (Fig. 2i, Supplementary Fig. 15a), and an MAE of 0.966% when mixing the near zero tumor fraction sample with the high tumor content sample (Supplementary Fig. 15b). Mixing bulk tissue samples at 30X target coverage resulted in slightly larger errors, with an MAE of 0.116% for the mixture containing the low tumor fraction sample (Supplementary Fig. 15c), while the mixture containing the high tumor fraction sample had an MAE of 1.60% (Supplementary Fig. 15d). Overall, cfClone produced consistent results for mixtures derived from either cfDNA or bulk tissue, suggesting that the model is not significantly confounded by cfDNA specific biases. We observe that in the high tumor fraction setting, cfClone infers estimates of tumor fraction that are in line with the predicted theoretical tumor fraction, while slightly underestimating tumor fraction in mixtures with very low tumor content.

### Real data validation

To assess the performance of cfClone with real cfDNA data, we compared cfClone to CloneSeq-SV, a scWGS informed panel based sequencing strategy designed for clonal tracking^12^. Both cfClone and CloneSeq-SV employ scWGS of patient tumors to identify cancer clones on the basis of HSCNVs. To deconvolute the clonal contributions to cfDNA, CloneSeq-SV employs targeted sequencing of clone-specific SNVs or SVs in contrast to cfClone’s whole genome approach.

We first compared the estimated tumour fraction estimates from cfClone with estimates obtained from error corrected duplex sequencing of clonal TP53 mutations (Spearman’s ρ = 0.85, p < 0.01; Fig. 3a). Similar results were obtained when comparing cfClone TF estimates to those obtained from using panels of either clonal SVs (Spearman’s ρ = 0.93, p < 0.01; Supplementary Fig. 16a) or clonal SNVs (Spearman’s ρ = 0.96, p < 0.01; Supplementary Fig. 16b).

**Fig. 3.**
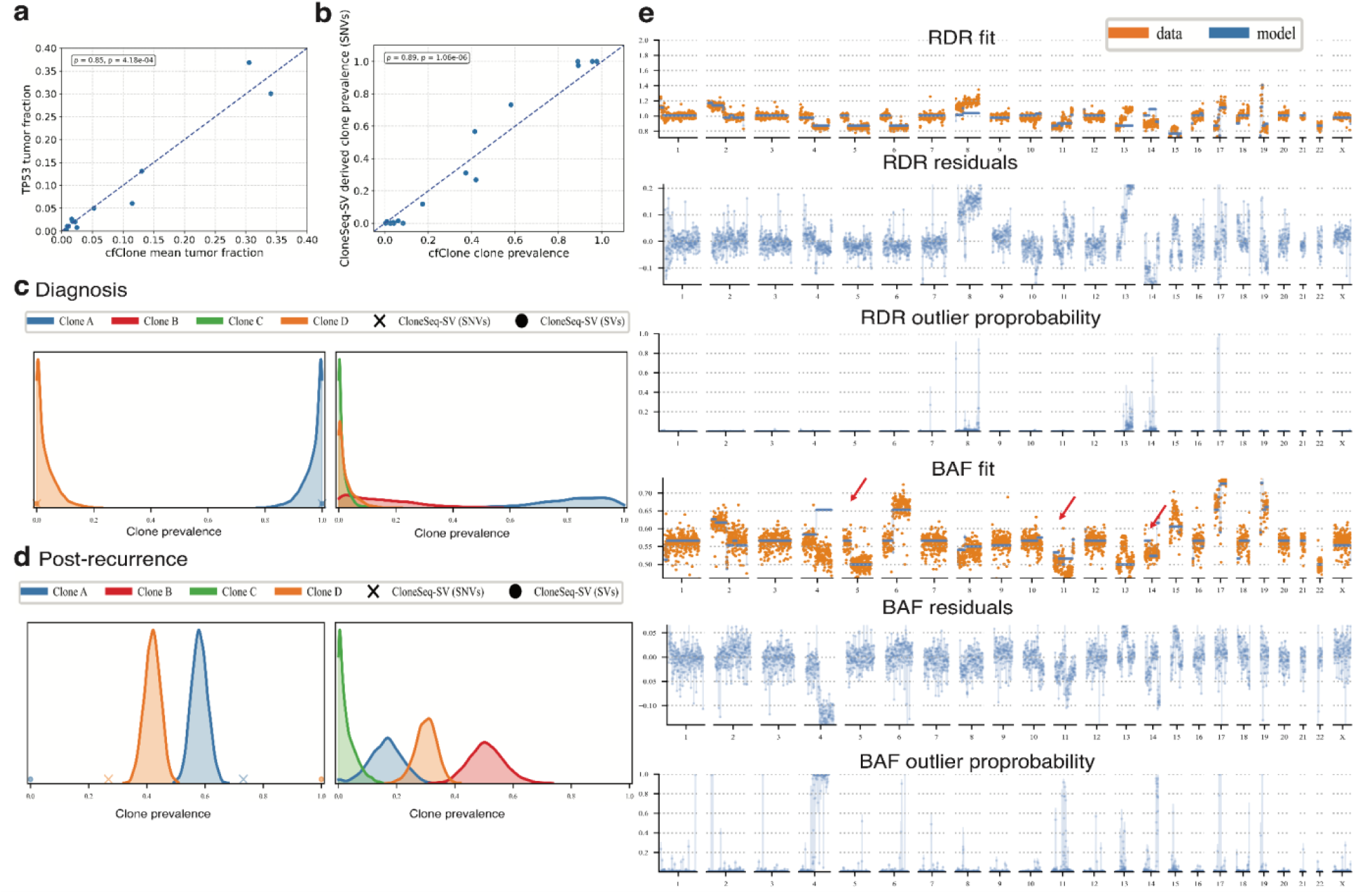
Validation on the MSK-SPECTRUM dataset. **a**, The mean tumor fraction per sample estimated by cfClone versus the tumor fraction estimated from *TP53* VAF (n = 6 patients, 12 samples). **b**, The mean clonal prevalence estimated by cfClone versus the clonal prevalence estimated by CloneSeq-SV using SVs (n = 3 patients, 6 samples). **c**, The posterior density of clonal prevalence predicted by cfClone for patient SPECTRUM-OV-107 at diagnosis (timepoint 1) under the two-clone (left) and four-clone (right) models. Density was smoothed with kernel density estimation. **d**, The posterior density of clonal prevalence predicted by cfClone for SPECTRUM-OV-107 after recurrence (timepoint 5) under the two-clone (left) and four-clone (right) models. Density was smoothed with kernel density estimation **e**, Model fits for patient SPECTRUM-OV-044 5 days after the time of first surgery (timepoint 1); fits are shown in RDR-space and BAF-space, along with their corresponding residuals and outlier probabilities. Red arrows depict a selection of missed HSCN events.

We next assessed the accuracy of cfClone’s clonal prevalence estimates. Of the six cases with bulk WGS from the ClonSeq-SV publication, three had clone sets which CloneSeq-SV could distinguish using both SVs and SNVs. The estimates of clonal prevalence produced by cfClone for these three cases were significantly correlated with CloneSeq-SV estimates derived from either SVs (Spearman’s ρ = 0.76, p < 0.01; Fig. 3b) or SNVs (Spearman’s ρ = 0.89, p < 0.01; Supplementary Fig. 16c). Of note, cfClone’s concordance with CloneSeq-SV’s SV or SNV derived clonal prevalences is comparable to CloneSeq-SV’s internal concordance between its SV and SNV-derived clonal prevalences for these three cases (Spearman’s ρ = 0.83, p < 0.01; Supplementary Fig. 16d).

Next we explored how the previously published results varied when including the full set of HSCNV defined clones from scWGS in the cfDNA analysis. Owing to the inability to distinguish all clones present in a given case on the basis of SVs or SNVs, CloneSeq-SV could only operate on a subset of identified clones in some cases. As cfClone only requires HSCNV profiles for each clone, it can distinguish the full set of clones identified by scWGS in cfDNA samples.

The consequence of using a reduced clone set is illustrated by the case of SPECTRUM-OV-107 (Fig. 3c-d), which features four clones (A-D). In this case only two clones were uniquely identified on the basis of SVs or SNVs. Using the same two clones, denoted A and D, cfClone (Fig. 3c, left; mean clonal prevalence; A = 97.6%, 95% HDI [91.9%, 100.0%] ; D = 2.43%, 95% HDI [0.00%, 8.07%]) and CloneSeq-SV (Fig. 3c, left; mean clonal prevalence; A = 100%, D = 0.00%) produce similar estimates of clonal prevalence at diagnosis, with clone A initially accounting for almost all clonal prevalence. In contrast, running cfClone with all four clones that were resolvable by scWGS, produces a significant shift in the clonal proportions at diagnosis (Fig. 3c, right). Using the full clone set with cfClone revealed that while clone A is still the most prevalent (mean clonal prevalence = 66.5%, 95% HDI [48.1%, 83.9%]), clone B also contributed a significant proportion of ctDNA (mean clonal prevalence = 31.1%, 95% HDI [12.0%, 48.3%]). Bayesian model selection suggests that there is insufficient evidence to prefer the two- or four-clone model at diagnosis (BF = 0.01), but the four-clone model is strongly favoured at the post-recurrence time point (BF = 36).

In line with this observation, while both methods produce comparable estimates of clonal prevalence under the two-clone model at diagnosis, they differ significantly under the two-clone model after recurrence (Fig. 3d). Based on SVs, CloneSeq-SV assigns all clonal prevalence to clone D following recurrence, reflecting an inversion of clonal prevalence at diagnosis (Fig. 3d, left). In contrast cfClone places nearly equal prevalence on both clones, modestly favouring clone A (Fig. 3d, left; mean clonal prevalence; A = 58.1%, 95% HDI [52.7%, 63.6%]; D = 41.9%, 95% HDI [36.4%, 47.3%]). This discrepancy is readily accounted for by the fact that CloneSeq-SV’s estimate is based on the detection of only a single SV in clone D following recurrence (Supp. Fig. 16e). Indeed in the two-clone model, using SNVs rather than SVs, CloneSeq-SV predicts that clone A dominates clone D (Fig. 3d, left; mean clonal prevalence; A = 73.2%, D = 26.8%). Under the four-clone model, cfClone predicts that clone B is the dominant clone following recurrence, with significant additional contributions from clones A and D (Fig. 3d, right; mean clonal prevalence; A = 16.2%, 95% HDI [5.29%, 28.0%]; B = 50.9%, 95% HDI [39.0%, 62.8%]; C = 2.78%, 95% HDI [0.00%, 8.89%]; D = 30.1, 95% HDI [22.9%, 37.2%]).

Due to the use of panel sequencing CloneSeq-SV, cannot identify the presence of clones not detected by scWGS of tissue. In contrast, cfClone models HSCNVs across the entire genome and provides a fitted model which can be used to identify regions of poor fit (high residuals). Where these residuals are sufficiently large in a genomic region, as measured by the RDR/BAF outlier probability, we can infer that a novel clone defining HSCNV events in these regions are contributing ctDNA to the sample.

This is well illustrated by the case of SPECTRUM-OV-044^12^, which features four clones. The model BAF fit and resulting residuals (Fig. 3e) suggest that one or more clones with a large HSCNV on one arm of chromosome 4 was missed; similar smaller missed events can be seen on chromosomes 11 and 13 in BAF-space, along with a corresponding missed gain on chromosome 13 in RDR-space. Whether these missed events are attributable to a single or multiple clones cannot be determined by cfClone. Nevertheless cfClone provides visual and quantitative diagnostics to identify regions of poor fit, indicative of the presence of missed or emergent clones.

### BCCA cohort analysis

To further evaluate the performance of cfClone in a clinical setting, we analyzed three patient cases with a diagnosis of HGSOC. For each case, banked biospecimens were available, including serial blood samples collected at multiple time points and retrospective surgical fresh-frozen tumor tissue obtained from treatment at the BC Cancer Agency (BCCA). Each patient had at least one available surgical tumor specimen along with longitudinal blood samples collected at distinct time points. The first case (6608) represents a patient treated with carboplatin and paclitaxel following debulking surgery (Fig. 4a, Event track). Plasma obtained 2 days before surgery (day 32) had a mean tumor fraction of 1.32% (95% HDI [1.28%, 1.37%]; Fig. 4b, TF track), with positive ctDNA detection by Bayes factor test (see legend, ctDNA detection). At this timepoint a single clone, K, dominates, depicted visually by its large mean clonal prevalence (45.1%, 95% HDI [35.2%, 54.6%]; Fig. 4a, CP track) and by its dominance in the pairwise rank plot of the top clones (Fig. 4b), with minor contributions from clones D, E, H, and L. By the final timepoint following chemotherapy (day 269) tumor fraction remains detectable, rising to a mean of 3.04% (95% HDI [2.98%, 3.09%]). Accompanying the overall rise in tumor fraction, clonal diversity increased following treatment, with clone K falling to a mean prevalence of 19.9% (95% HDI [15.6%, 24.8%]), with a large rise in the prevalence of clones D and L, and inversion of the prevalence of clones E and H (Fig. 4a, CP track). Similarly in the pairwise rank plot (Fig. 4b), clone K ceases to be dominant, with clones D and L dominating clones K, E, and H, while clone E dominated H.

**Fig. 4.**
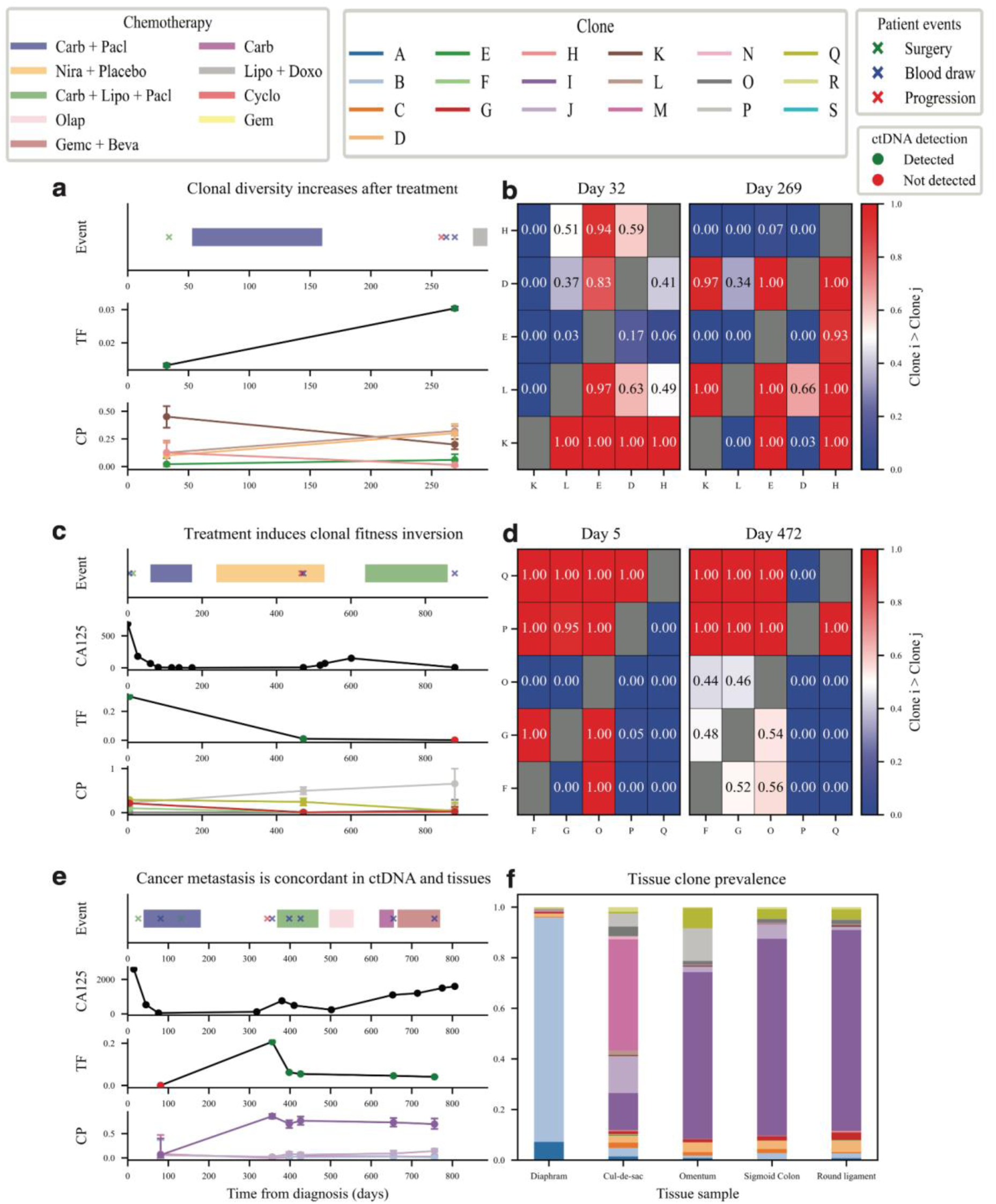
Application of cfClone to the BCCA cohort. **a**, Patient events (Event), mean tumor fraction (TF), and mean clonal prevalence (CP) plots for case 6608. **b**, Pairwise rank plots for case 6608; plots give the probability that clone i (row) is greater than clone j (column). **c**, Event, CA125, TF, and CP plots for case 6607. **d**, Pairwise rank plots for case 6607 at the first (day 43) and second (day 510) blood draw timepoints. **e**, Event, CA125, TF, and CP plots for case 6606. **f**, Mean clonal prevalence inferred from PDXs derived from various metastatic sites for case 6606.

In the second case (6607) a blood draw was obtained on day 5, 9 days before debulking surgery, with a high mean tumor fraction of 30.3% (95% HDI [30.2%, 30.4%]; Fig. 4c, TF track), corresponding with the highest CA125 value observed for this patient (687 U/ml; Fig. 4c, CA125 track). At this time the most abundant clones were G, F, O, P, and Q, with the greatest contribution from Q (Fig. 4c, CP track; Fig. 4c). Following surgery the patient was treated with carboplatin and paclitaxel (Fig. 4c, Event track), followed by a break before additional treatment with niraparib. During treatment with niraparib the patient progressed and a blood sample was obtained 5 days later (day 472, Fig. 4c, Event track). At this time the mean tumor fraction was 1.07% (95% HDI [1.03%, 1.11%]) with positive ctDNA detection (Fig. 4c, TF track), with a correspondingly low CA125 of 6.2 U/ml. An inversion in clonal fitness was apparent, with clone Q ceasing to dominate, having been overtaken in prevalence by clone P (Fig. 4c, CP track), suggesting that either more residual tumour remained containing clone P after surgery or that clone P was refractory to treatment; assessed by pairwise rank, P had become dominant over all clones, with Q remaining the second most dominant (Fig. 4d). The patient then received a final treatment with carboplatin, liposomal doxorubicin, and paclitaxel. The final blood sample obtained near the time of death did not contain detectable amounts of ctDNA (Fig. 4d, TF track).

Finally we examined a case (6606) for which 6 time-series blood samples were obtained throughout treatment, permitting longitudinal tracking of clonal prevalence. This patient harboured a germline TP53 mutation consistent with Li-Fraumeni syndrome and a somatic *BRCA1* mutation. They presented with metastatic disease and underwent primary debulking surgery followed by interval debulking surgery during initial treatment with carboplatin and paclitaxel (Fig. 4e, Event track). Blood draws during the first round of chemotherapy contained no detectable ctDNA (Fig. 4e, TF track) along with a modest CA125 of 50 U/ml (Fig 4e, CA125 track). Following the first round of treatment the patient progressed; a blood draw shortly after showed that the mean tumor fraction had risen to 20.7% (95% HDI [20.6%, 20.8%]; Fig. 4e, TF track), with a substantial increase in CA125 to 760 U/ml 24 days later (Fig. 4e, CA125 track). Just after progression, clone I was assessed to be significantly more abundant than all other clones, with a mean clonal prevalence of 70.2% (95% HDI [61.6%, 77.5%]; Fig. 4e, CP track). The patient treatment history included first-line carboplatin plus paclitaxel (6 cycles total), second-line carboplatin plus liposomal doxorubicin followed by maintenance olaparib, third-line carboplatin monotherapy, and fourth-line gemcitabine plus bevacizumab. Tumor fraction was initially reduced during treatment to 6.19% (95% HDI [6.13%, 6.24%]), but remained stable and detectable throughout the remaining treatments (Fig. 4e, TF track). Similarly, clone I remained the dominant clone throughout treatment (Fig. 4e, CP track). In contrast with tumor fraction, CA125 rose continuously throughout treatment, peaking at 1600 U/ml in the final timepoint (Fig. 4e, CA125 track). In this case, WGS of multiple metastases enabled us to assess the concordance between the clonal prevalence observed in blood and that present at metastatic sites. Bulk WGS data was generated from tumor samples obtained from the diaphragm, cul-de-sac, omentum, sigmoid colon, and round ligament (Fig. 4f). We applied cfClone to deconvolve the clonal proportions present at each site. There was broad concordance between the dominant clone observed in ctDNA and the peritoneal metastatic sites, with clone I accounting for the vast majority of clonal prevalence in the omentum (66.0%, 95% HDI [64.2%, 67.7%]), sigmoid colon (78.0%, 95% HDI [76.2%, 79.6%]), and round ligament (79.5%, 95% HDI [77.6%, 81.2%]; Fig. 4f); clones B and M were the dominant clones in the diaphragm and cul-de-sac, respectively, but only made minor contributions to ctDNA at any time (Fig. 4e, CP plot). This suggests that clone I, which was refractory to treatment, likely seeded many of the metastases, and that cfClone is readily applicable to the deconvolution of WGS obtained from bulk tumor samples in addition to cfDNA.

## Discussion

While previous studies have demonstrated the feasibility of reconstructing tumor clonal architecture from cfDNA, accurate longitudinal tracking of clonal dynamics remains challenging, particularly at low tumor fractions and sequencing coverage^16,10,12,17,18^. To address these limitations, we developed cfClone, a statistical method for high-resolution reconstruction and minimally invasive longitudinal tracking of tumor clonal architecture using a combination of scWGS and cfDNA. Using in-silico data derived from FL, DLBCL, and HGSOC, we demonstrate that cfClone provides more accurate estimate of tumor fraction compared to ichorCNA, reliably detecting tumor fractions as low as 0.25% at 100X coverage, with approximately an order of magnitude lower error across various combinations of coverage and tumor fraction. The use of a Bayesian approach enables cfClone to provide quantitative estimates of tumor fraction and clonal prevalence which incorporate uncertainty, and to robustly determine the presence/absence of ctDNA via a Bayes factor test.

Comparison with CloneSeq-SV, the only alternative method for deconvoluting clonal prevalence in cfDNA samples using matched scWGS, demonstrates that cfClone provides comparable estimates of tumor fraction and superior clonal tracking. Importantly, cfClone has no requirement for a bespoke panel of SVs/SNVs per patient, which is required by CloneSeq-SV. The use of clone HSCNV profiles enables cfClone to track all the clones detected from scWGS, including those which lack the distinguishing SVs or SNVs necessary for panel-based methods. In addition, cfClone provides metrics enabling the detection of clones which were missed in the paired scWGS sample, a feature absent from panel-based approaches but which is essential for a longitudinal context in which new clones may emerge following initial scWGS sampling. Although cfClone provides a means of identifying the presence of novel clones, future work will be required to determine the number and genotypes of novel clones.

Using real data we demonstrate time-series tracking of clonal prevalence in 3 HGSOC patients. Across these patients we observe highly divergent patterns in clonal prevalence in response to treatment, including an increase in clonal diversity, suggestive of polyclonal selection; an inversion in clonal prevalence; and longitudinal tracking of a single clone which became dominant in response to initial treatment with the standard of care (carboplatin and paclitaxel) and which thereafter demonstrated resistance to multiple therapies. In the latter case we show that cfClone is readily applicable to bulk tumor samples, and that the dominant clone in cfDNA following progression can be traced to specific metastatic sites.

We expect that cfClone’s capacity to quantify and track clones is broadly generalizable to genomically unstable cancers which exhibit significant aneuploidy, such as HGSOC, p53-abnormal endometrial cancer, triple-negative breast cancer (TNBC), and lung adenocarcinoma (LUAD). For applications limited to TF quantification and ctDNA detection, cfClone is readily applicable to cancers with limited CNV burden. This is exemplified in the FL-DLBCL mixtures as two clones from the DLBCL cancer type share many copy number events, which interfered with estimation of clonal prevalence, but cfClone still robustly estimated tumor fraction at low coverage and expected tumour content.

In summary we have developed cfClone, a Bayesian method for scWGS informed analysis of liquid biopsies. Through the analysis of synthetic and real world HGSOC data, we demonstrate that cfClone provides sensitive and accurate detection of ctDNA, which can be used for the detection of minimal residual disease. Furthermore, cfClone provides quantitative estimates of TF which include uncertainty, making it suitable for quantitative assessment of treatment response. Finally, cfClone decomposes TF into clonal prevalence estimates which can be used to determine clone specific response to therapy. This last point creates an opportunity to precisely assess the clone specific features driving differential response to treatment and personalise treatment strategies.

## Methods

### Clinical cohort description and ethical considerations

We analyzed cell-free DNA (cfDNA) samples from a retrospective cohort of 3 patients with high-grade serous ovarian cancer (HGSOC) enrolled in the GTB cohort between 2016 and 2023 in Vancouver. Vancouver patients underwent primary treatment at BC Cancer Agency (Vancouver, Canada). Ovarian cancer patients were recruited under Institutional Review Board approval from the University of British Columbia (BC Cancer REB approval H25-02463).

### DNA extraction from plasma and library prep

Plasma cfDNA libraries were prepared using the KAPA HyperPrep Kit (Roche) following an optimized protocol adapted from the New York Genome Center. Briefly, 10 ng of cfDNA input was subjected to end repair and A-tailing (20 °C for 30 min; 65 °C for 30 min), followed by ligation of diluted TruSeq DNA UD adapters (IDT) at 20 °C for 30 min. Adapter-ligated DNA was purified using SPRIselect magnetic beads (Beckman Coulter).

Libraries were amplified using KAPA HiFi HotStart ReadyMix with 6–8 PCR cycles, depending on input DNA quantity (< 5 ng: 8 cycles; > 5 ng: 6–7 cycles), under standard thermocycling conditions. Post-amplification cleanup was performed using SPRIselect beads. Final libraries were eluted in EB buffer (Qiagen) and quantified using Qubit high-sensitivity assays, with fragment size distribution assessed by Bioanalyzer. The expected library size was around 300 bp.

### Single-cell whole-genome sequencing and library construction of tumors with DLP+

Tumor tissues were dissociated into single-cell suspensions as previously described^2,19,20^. Briefly, PDX tumor fragments were incubated with collagenase/hyaluronidase, 1:10 (10X) enzyme mix (STEM CELL technologies, Catalog # 07912) in DMEM/F-12 with Glucose, L-Glutamine and HEPES (Lonza 12-719F) and 1% BSA (Sigma) at 37°C. After enzymatic digestion the final single cell pellet was resuspended in 0.04% BSA (Sigma) and PBS to achieve ∼1 million cells/ml concentration for robotic cell spotting onto the DLP+ chip.

### Robot spotting of single cells into the nanolitre wells and library construction

The construction of the DLP+ library was carried out as described in^2,19,20^. Briefly, single cell suspensions from patient derived xenografts were fluorescently stained using CellTrace CFSE (Life Technologies) and fixable Red Dead Cell Stain (ThermoFisher) LIVE/DEAD (ThermoFisher) in a PBS solution containing 0.04% BSA (Miltenyi Biotec 130-091-376) incubated at 37°C for 20 minutes. The cells were then centrifuged to remove stain and resuspended in fresh PBS with 0.04% BSA. This single cell suspension was loaded into a contactless piezoelectric dispenser (sciFLEXARRAYER S3, Scienion) and spotted in open nanowell arrays (SmartChip, TakaraBio) preprinted with unique dual index sequencing primer pairs. Libraries were sequenced at UBC Biomedical Research Centre (BRC) in Vancouver, British Columbia on the Illumina NextSeq 550 (mid- or high-output, paired-end 150-bp reads), or at the GSC on Illumina HiSeq2500 (paired-end 125-bp reads) and Illumina HiSeqX (paired-end 150-bp reads). The data was then processed via a quantification and statistical analysis pipeline^2,19,20^.

### Pre-processing of whole-genome sequencing data

Raw whole-genome sequencing (WGS) reads from bulk tumors, blood buffy coat (BC), and blood plasma cell-free DNA (cfDNA) samples were preprocessed using the same workflow. Per-sample reads were aligned separately for each read group (flow cell and lane) to the GRCh38 reference genome^21^ using bwa-mem v0.7.19. Following alignment, duplicate reads were marked with Genome Analysis Toolkit (GATK) v4.6.2.0^22^MarkDuplicatesSpark and base quality scores were recalibrated with BaseRecalibrator. Phasing was obtained from buffy coats via Eagle v2.4.1. For plasma cfDNA and bulk tumor samples, reads were counted in 500 kb bins and the ratio between the read depth in each bin and read depth in the respective normal sample was computed, yielding a RDR; RDR was then corrected for biases arising from GC-content and mappability using modal quantile regression. For cfDNA, a normal plasma sample with no evidence of disease (NED) was used; for tissues the patient’s buffy coat was used. BAF was computed for plasma cfDNA and bulk tumor samples by counting reads for each haplotype block.

### Pre-processing of single-cell whole-genome sequencing data

Single-cell whole-genome sequencing (scWGS) reads were preprocessed using an in-house pipeline. Within a sample, reads for each cell were aligned per read-group to the GRCh38 reference genome^21^ using bwa-mem v0.7.19. Duplicate reads were marked with GATK v4.6.2.0^22^. MarkDuplicatesSpark and base quality scores were recalibrated with BaseRecalibrator. Poor quality cells, such as those with low read depth were filtered out and the resulting per-cell BAM files were collected. Allele counts for each cell BAM were computed for a list of known SNPs and phased with Eagle v2.4.1^23^. Reads in each cell BAM were counted into 500 kb bins, corrected for GC-content and mappability biases. For each patient, corrected read data and haplotype block counts were provided as input to our in-house CNV calling tool HapClone v0.1.0, yielding haplotype-specific clone copy-number profiles.

### Processing of MSK-SPECTRUM data

Pre-processed data for the MSK-SPECTRUM cases was obtained from the supplementary information of the associated paper^12^, along with the Synapse repository (syn66399325) and processed with the spectrum-ctdna pipeline. The RDR and BAF data for each cfDNA timepoint per patient, along with the corresponding clone copy-number profiles, were provided as input to cfClone v0.3.1. For each patient cfClone was run with the full set of clones from the respective patient, as well as the subset of patient clones which were uniquely identifiable via SVs and/or SNVs in William et al.^12^. All analyses of estimated mean tumor fraction and mean clonal prevalence are derived from the final model restart, numbered ‘2’.

### Processing of the BCCA patient cohort data

Following pre-processing of the bulk and single-cell whole-genome sequencing data, the clone HSCN profiles for each patient derived from HapClone v0.1.0 were provided as input to cfClone v0.3.1 along with the patient’s respective plasma cfDNA and/or bulk tumor RDR/BAF profiles. All analyses of estimated mean tumor fraction and mean clonal prevalence are derived from the first model restart, numbered ‘0’.

## Supporting information

https://drive.google.com/file/d/1gW3PQp38THNQyM-QOtpm8o9LHb4SPosT/view?usp=drive_link

## Acknowledgements

This work was supported by the ERC Synergy grant (LUMES, Grant #101224965) and Terry Fox Research Institute (TFRI Project Grant #1162-06). A.R. is supported by the BC Cancer Foundation and the Natural Sciences and Engineering Research Council of Canada (NSERC; Grant RGPIN-2022-04378). E.H. is supported by a Canadian Institutes of Health Research (CIHR) Canada Graduate Scholarship–Doctoral (CGS-D) (Award #209937). F.K. is supported by a Michael Smith Health Research BC Research Trainee Award (Grant #RT-2023-3343) and a Canadian Institutes of Health Research (CIHR) Postdoctoral Fellowship (Award #202110MFE-472574-205950).

## Code and data availability

Code for cfClone can be found at https://github.com/Roth-Lab/cfclone. Raw WGS data from cfDNA, buffy coats, and bulk tumors will be deposited at the European Genome-phenome Archive (EGA) upon acceptance. Raw scWGS data was previously published and is available through the EGA, accession number EGAS00001006343. For schematic figures biorender was used under individual academic agreement number XS2A0KAJ3P to Kabeer, F.

