## Supplementary material for "Minimally invasive monitoring of clonal evolution through integrated single cell and ctDNA analysis": https://drive.google.com/file/d/1gW3PQp38THNQyM-QOtpm8o9LHb4SPosT/view?usp=drive_link

### 1 Supplementary Methods: cfClone model

We describe in this section the technical underpinnings of cfClone. Recall that at a high level, cfClone models cfDNA as a mixture of DNA fragments originating from $K$ distinct cancer cell populations and the normal cell population. Each of the $K$ populations is characterized with its copy number profile, which is obtained from single cell whole genome sequencing (scWGS) of a matched tissue sample as a pre-processing step.

The primary parameter of interest is the clonal prevalence probability vector (also known as simplex) $\rho$ which quantifies the contribution of both the normal population as well as each cancer cell population. Specifically, $\rho = (\rho_1, \ldots, \rho_K, \rho_{K+1})$ denotes a simplex-valued parameter of size $K+1$, $\sum_{k=1}^{K+1} \rho_k = 1$, where the last component, $n = K+1$, is reserved for the normal cell prevalence. We interpret each component $\rho_k$ as the fraction of cells shedding into the blood stream that belong to population $k$. Consequently, we can estimate the overall tumour fraction as a by-product, by summing over the cancer clone prevalences, $\sum_{k=1}^{K} \rho_k$.

#### 1.1 Probability model

The cfClone model utilizes two genomic data modalities: corrected read depth ratios (RDR) and B-allele frequency (BAF), each defined in more detail below. Each modality is considered conditionally independent given the clonal proportions and clone copy number profiles. The prevalence parameter $\rho$ is inferred using RDR and BAF data jointly, allowing cfClone to take into account both total and haplotype-specific copy number variations.

In the following, we use $t$ to index over genomic bins, and we let $T$ denote the total number of genomic bins, $t \in [T]$. Let $C_{tk}$ denote the copy number of population $k \in [K+1]$ at genomic bin $t$, inferred from scWGS matched tissue. We also assume that haplotype phasing has been performed as pre-processing for each population, yielding a decomposition of the total copy number into haplotype copy numbers $C^a_{tk}, C^b_{tk}$ satisfying $C_{tk} = C^a_{tk} + C^b_{tk}$. The preprocessed haplotype copy numbers are also derived from scWGS matched tissue.

A central component of cfClone is a Bayesian model, i.e., a joint probability distribution over the main parameter of interest, $\rho$, the observations, and nuisance parameters. The remainder of this section defines formally that joint probability distribution. We first describe each of the two likelihood models, one for RDR and one for BAF, and conclude this section with the specification of the priors.

**RDR likelihood model.** There are two sources of data needed for the RDR likelihood model. First, the liquid biopsy data itself, with sufficient statistics denoted as follows: $T_t$ for the number of reads in bin $t$ in the liquid biopsy, and $N$, for the total number of reads, $N = \sum_t T_t$. Second, a matched baseline cfDNA, which can come from a non-evaluable disease (NED) sample, or, if a NED sample is not available, a surrogate such as a buffy coat could be considered as an inferior alternative. We use similar notation: let $\check{T}_t$ denote the number of reads in bin $t$ in the baseline cfDNA, and $\check{N}$, the corresponding total number of reads, $\check{N} = \sum_t \check{T}_t$. Following previous work[1], from these two sources of data, define the RDR as

$$L_t = \frac{(T_t/N)}{(\check{T}_t/\check{N})}. \tag{1}$$

The relationship between the observed RDR data and latent clonal contributions is captured through deterministic functions denoted by $\mu_t$, which take hypothesized values of $\rho$ as input. More precisely, in Section 1.2, we show that under reasonable assumptions we have

$$L_t \approx \mu_t := \alpha \times \frac{\sum_k \rho_k C_{tk}}{\frac{1}{T}\sum_t \sum_k \rho_k C_{tk}}, \tag{2}$$

where from now on, "$\sum_k$" denotes "$\sum_{k=1}^{K+1}$", i.e., the normal population is included in the summations over $k$ unless specified otherwise, and $\alpha$ is a batch-specific positive constant. In our model, we learn the nuisance parameter $\alpha$ jointly. Motivated by Equation (2), the core of the RDR likelihood model is a student-$t$ distribution with location parameter $\mu_t$, scale $\sigma$ and degrees of freedom $\nu$, denoted Student-$t(\nu, \mu_t, \sigma)$. We learn the parameter $\sigma$ jointly, and fix $\nu = 25$. Finally, to help visualize the effect of missing and/or emergent clones, we construct a mixture model for each genomic site $t$, where the first component is the informed distribution that we have just defined, Student-$t(\nu, \mu_t, \sigma)$, and the second component is a diffuse distribution, namely Student-$t(4, 1, 1)$. Each genomic bin has a distinct RDR outlier indicator variable, $z_t^{\mathrm{rdr}} \sim \mathrm{Bernoulli}(\pi_{\mathrm{rdr}})$. All indicator variables $z_t^{\mathrm{rdr}}$ are analytically marginalized during MCMC sampling. We also developed a post-processing step re-instantiating the posterior mean of $z^{\mathrm{rdr}} = (z_1^{\mathrm{rdr}}, \dots, z_T^{\mathrm{rdr}})$ to assist residual analysis when searching for missing/emergent clones. To summarize, the RDR likelihood is defined by:

$$L_t \sim \pi_{\mathrm{rdr}} \cdot \text{Student-}t(4, 1, 1) + (1 - \pi_{\mathrm{rdr}}) \cdot \text{Student-}t(25, \mu_t, \sigma),$$

where the RDR outlier mixing proportion, $\pi_{\mathrm{rdr}}$, which is shared by all RDR outlier indicator variables, is learned jointly.

**BAF likelihood model.** Let $D_t$ denote the number of read counts in the liquid biopsy that overlap heterozygous SNPs in genomic bin $t$. Let $B_t \in \{0, 1, \dots, D_t\}$ denote how many of those match with haplotype B (recall that we assume haplotyping of the scWGS data has been performed as a preprocessing step). In brief, the construction of the BAF likelihood model follows similar steps as for its RDR counterpart: we first construct a location parameter (in the BAF case, a mean frequency $p_t$ instead of $\mu_t$ for RDR), then we build a distribution centered at that mean (in the BAF case, a BetaBinomial, but with an alternative parameterization compared to the standard BetaBinomial), and finally, we use a mixture to help detection of missing/emergent clones. Next, we go over each of these three steps in more detail. For the first step, the predicted frequency of haplotype B in bin $t$ is simply given by:

$$p_t := \frac{\sum_k \rho_k C_{tk}^b}{\sum_k \rho_k C_{tk}}.$$

For the second step, we describe first the model using a Beta nuisance variable to help explain the change of parametrization, but that nuisance variable is analytically marginalized in the implementation. We let $B_t \sim \mathrm{Binomial}(\tilde{B}_t, D_t)$, and place a Beta distribution on $\tilde{B}_t$. The classical parameterization for the Beta family, in terms of shape parameters $\alpha, \beta > 0$, is not straightforward to inform by $p_t \in [0, 1]$, so a common alternative is the Beta mean-concentration parameterization, $p \in [0, 1], \kappa > 0$, given by $\alpha = p\kappa$, $\beta = (1-p)\kappa$, for which $p$ is the mean. However, we found that the parameter $\kappa$ concentrates on large values, so we introduce a novel reparametrization, $\gamma := 1/\kappa \in (0, 1)$. Since $\gamma \to 0$ leads to a straightforward binomial likelihood, the parameter $\gamma$ has an intuitive interpretation as controlling "divergence from binomiality" (abbreviated db). After marginalization of $\tilde{B}_t$, we denote the obtained parameterization by $\mathrm{BetaBinomial}_{\mathrm{db}}(D_t, p_t, \gamma)$. Moving on to the mixture step, the outlier component is this time given in the standard $(\alpha, \beta)$ parameterization by $\mathrm{BetaBinomial}(D_t, \alpha = 1, \beta = 1)$. Denote the BAF outlier indicator variables by $z_t^{\mathrm{baf}} \sim \mathrm{Bernoulli}(\pi_{\mathrm{baf}})$, which again are marginalized during MCMC sampling but re-instantiated for post-processing. To summarize, the BAF likelihood is defined by:

$$B_t \sim \pi_{\mathrm{baf}} \cdot \mathrm{BetaBinomial}(D_t, \alpha = 1, \beta = 1) + (1 - \pi_{\mathrm{baf}}) \cdot \mathrm{BetaBinomial}_{\mathrm{db}}(D_t, p_t, \gamma),$$

where the BAF outlier mixing proportion, $\pi_{\mathrm{baf}}$, which is shared by all BAF outlier indicator variables, is learned jointly. Note that in contrast to the RDR likelihood model, the BAF likelihood model does not depend on the baseline cfDNA data.

**Prior distributions.** As customary for simplex-valued variables, we use a Dirichlet prior on $\rho$, defined with the usual parameterization $(\alpha_1, \alpha_2, \dots, \alpha_K, \alpha_{K+1})$, $\alpha_k > 0$. We use a hyper-parameter of $\alpha_k = 1/2$ for the clones, $k \in [K]$ and $\alpha_{K+1} = 10$ for the normal population. Higher value on the normal population reflects the prior belief that most DNA fragments in a cfDNA sample will be derived from healthy cells. The lower values (1/2) for the clonal prevalences act as a regularizer encouraging near-sparsity unless dictated otherwise by the data. Values lower than 1/2 can lead to numerical issues. In the following, $\mathrm{Gamma}(\alpha, \beta)$ always refers to the shape-rate parametrization (large values of $\beta$ lead to concentrated priors). For the shift parameter $\alpha$, we use a $\mathrm{Gamma}(1, 1)$, a distribution with a mean of one, motivated by the analysis in Section 1.2. For the scale parameter $\sigma$ used for the RDR likelihood, we use a prior concentrated on small values, namely Gamma(1, 100). The last three parameters, $\pi_{\text{rdr}}$, $\pi_{\text{baf}}$ and $\gamma$ are also expected to be small, but this time with range $[0, 1]$, so we use Beta($\alpha = 1, \beta = 100$) priors for those parameters. To summarize:

$$\begin{aligned}
\rho &\sim \text{Dirichlet}(\underbrace{1/2, 1/2, \ldots, 1/2}_{\text{for the } K \text{ clones}}, 10) && \text{(population prevalences)} \\
\alpha &\sim \text{Gamma}(1, 1) && \text{(shift term used in the RDR likelihood)} \\
\sigma &\sim \text{Gamma}(1, 100) && \text{(Student-}t\text{ scale parameter used in the RDR likelihood)} \\
\pi_{\text{rdr}} &\sim \text{Beta}(1, 100) && \text{(outlier rate for RDR bins)} \\
\pi_{\text{baf}} &\sim \text{Beta}(1, 100) && \text{(outlier rate for BAF bins)} \\
\gamma &\sim \text{Beta}(1, 100) && \text{(divergence from binomiality parameter, BAF likelihood).}
\end{aligned}$$

To avoid numerical issues in our implementation, we replaced all parameter constraints of the form $x > 0$ by $x > 10^{-6}$.

#### 1.2 Theoretical analysis supporting RDR likelihood model construction

In this section we prove a theoretical result supporting the functional form we use for the location parameter of the RDR likelihood model, namely:

$$\mu_t = \alpha \times \frac{\sum_k \rho_k C_{tk}}{\frac{1}{T} \sum_t \sum_k \rho_k C_{tk}}.$$

To state the result, consider a baseline cfDNA obtained from a diploid genome, and let $\xi_t \in [0, 1]$ denote the probability that one read in the baseline cfDNA falls in bin $t$. Denote the vector of these probabilities by $\xi = (\xi_1, \xi_2, \ldots, \xi_T)$, $\sum_t \xi_t = 1$. Even for a diploid genome, $\xi$ will deviate from the uniform distribution, $\xi \neq \xi_{\text{unif}} = (1/T, 1/T, \ldots, 1/T)$. This is due to a range of biological and technical reasons, including the binding of DNA to histones, GC bias and non-uniform mappability[2–4].

Recall that the RDR is computed[1] as

$$L_t = \frac{(T_t/N)}{(\check{T}_t/\check{N})}, \tag{3}$$

where $T_t$ is the number of reads in bin $t$ in the liquid biopsy, $N$ is the total number of reads, $N = \sum_t T_t$, and $\check{T}_t, \check{N}$ are the corresponding baseline cfDNA derived quantities. The main result of this section studies $L_t$ in the regime of a large number of total reads for both the liquid biopsy and baseline cfDNA:

**Proposition 1.** *If $N$ and $\check{N}$ go to infinity, we have:*

$$L_t \to \mu_t = \alpha \times \frac{\sum_k \rho_k C_{tk}}{\frac{1}{T} \sum_t \sum_k \rho_k C_{tk}}, \text{ almost surely,}$$

*with $\alpha = 1$ when $\xi = \xi_{unif}$, otherwise, $\alpha$ is some constant that does not depend on $t$.*

We start by applying the law of large numbers to each of the numerator and denominator in the definition of $L_t$. For the denominator, we have that for $\check{N}$ large, we expect $\check{T}_t/\check{N} \approx \xi_t$. For the numerator, $T_t/N$, we make the assumption that for the liquid biopsy sample, the probability for one read to fall in bin $t$ is proportional to $\xi_t \sum_k \rho_k C_{tk}$. Combining this assumption with the law of large numbers, we obtain that for $N$ large, we expect $T_t/N \approx F_t^\xi$, where

$$F_t^\xi = \frac{\xi_t \sum_k \rho_k C_{tk}}{\sum_{t'} \xi_{t'} \sum_{k'} \rho_{k'} C_{t'k'}}.$$

To analyze $F_t^\xi$, we will use the following lemma:

**Lemma 1.** *Let $F_t := F_t^{\xi_{unif}}$. We have $F_t^\xi = \xi_t F_t C_\xi$, for a positive constant $C_\xi$, i.e., $C_\xi$ does not depend on $t$ (but it depends on $\xi$, $\rho$ and on the copy number profiles). Moreover, if $\xi_t = \xi_{unif}$, we have $C_\xi = T$.*

*Proof.*

$$\frac{F_t^\xi}{\xi_t F_t} = \frac{\left(\xi_t \frac{\sum_k \rho_k C_{tk}}{\sum_{t'} \xi_{t'} \sum_{k'} \rho_{k'} C_{t'k'}}\right)}{\xi_t \left(\frac{\sum_k \rho_k C_{tk}}{\sum_{t'} \sum_{k'} \rho_{k'} C_{t'k'}}\right)}$$
$$= \frac{\sum_{t'} \sum_{k'} \rho_{k'} C_{t'k'}}{\sum_{t'} \xi_{t'} \sum_{k'} \rho_{k'} C_{t'k'}} =: C_\xi.$$

From the expression for $C_\xi$ given above, we see that if $\xi_t = \xi_{\text{unif}}$, then $C_\xi = T$. □

We can now complete the proof of Proposition 1:

*Proof.* Applying the law of large numbers twice and combining the ratios using the continuous mapping theorem, we obtain

$$L_t = \frac{(T_t/N)}{(\check{T}_t/\check{T})}$$
$$\to \frac{F_t^\xi}{\xi_t}, \text{almost surely.}$$

Next, applying Lemma 1, we have:

$$\frac{F_t^\xi}{\xi_t} = \frac{\xi_t F_t C_\xi}{\xi_t}$$
$$= F_t C_\xi.$$

Finally, from Lemma 1, we have that $\xi = \xi_{\text{unif}}$ implies $C_\xi = T$, and hence $\alpha = 1$. □

Going back to the full Bayesian inference context, the fact that $C_\xi$ and hence $\alpha$ depend on $\rho$ is not a problem. Indeed, since we approximate a full posterior distribution over both, we can learn non-linear relationships between the unknown variables, encoded as a correlated posterior distribution. Moreover, as we cover shortly, the MCMC methods we use are well equipped to efficiently explore correlated posterior distributions. In that context, the main result in this section states that the MCMC sampler can find some $\alpha$ and $\rho$ that will explain the RDR data well thanks to the form of $\mu_t$ that we use.

### 2 Residual analysis to reveal missing or emergent clones

A residual analysis on cfClone's model fit can be used as a qualitative diagnostic to identify missing or emerging clones. Suppose that an existing clone does not have an accompanying clone copy number profile inputted into cfClone by $C^a_{tk}$, $C^b_{tk}$. This would lead to a poor model fit as cfClone will not be able to model the copy number events that characterize the missing clone. There are two possible scenerios in which there is a missing clone. First, the clone cell population was not physically sampled during the patient's initial debulking surgery, and therefore, missed in the scWGS data. This would lead to the inference procedure using scWGS as input to be ignorant of the present clone. Additionally, it is also possible for the inference results to miss the clone despite the clone being present in the scWGS data such as when an initial clone cell population is small relative to other clones. Secondly, the cancer progresses and a clone emerges that was not present initial in the debulking surgery, and therefore, not present in the scWGS data. It is important to note that this generalizes to more than one clone and the residuals would be interpreted as the mixture of all the mixing clones.

#### 2.1 Summarizing the cfClone posterior distribution for cfDNA analysis

In this section, we first describe how we summarize the cfClone posterior distribution for cfDNA analysis, and then describe how we formulate the task of ctDNA detection as Bayesian model selection. We delay information on how the samples and Bayes factors are computed to Section 2.2.

**Posterior summaries.** The output of the MCMC sampler (Section 2.2) is a collection of $M$ post warm-up samples, denoted

$$\{(\rho^{(m)}, \alpha^{(m)}, \sigma^{(m)}, \pi_{\text{rdr}}^{(m)}, \pi_{\text{baf}}^{(m)}, \gamma^{(m)})\}_{m=1}^{M}.$$

Posterior means are approximated by averaging over the MCMC samples. For example, to compute a point estimate for the prevalence of population $k$, $\hat{\rho}_k$, we use

$$\hat{\rho}_k = \frac{1}{M}\sum_{m=1}^{M} \rho_k^{(m)}.$$

For a point estimate of the overall tumour fraction $\rho_{\text{f}}$, we use the complement of the normal population point estimate, indexed $n = K+1$, i.e., $\hat{\rho}_{\text{f}} = 1 - \hat{\rho}_n$.

We add uncertainty quantification around the point estimates using credible intervals. We use Highest Density Intervals (HDI) computed using the `ArviZ` package[5]. We use a nominal coverage of 95% unless specified otherwise.

To better understand the relationship between relative clone prevalences $\rho$ and the uncertainty of their estimates, we report, for all $k, k' \in [K]$ the Monte Carlo approximation of the posterior probability $\mathbb{P}(\rho_k > \rho_{k'}|\text{data})$:

$$d_{k,k'} = \frac{1}{M}\sum_{m=1}^{M} \mathbb{1}[\rho_k^{(m)} > \rho_{k'}^{(m)}],$$

which has the interpretation of being our belief given the data that clone $k$ has a larger prevalence than that of clone $k'$. For any $k, k'$, the above equation is the Bayes estimator corresponding to the action of deciding the ranking of $k$ and $k'$ with a 0-1 loss, and as such the estimator benefits from the Bayes estimator's optimality properties[6]. In particular, it is superior to heuristic criteria based on credible interval overlap. This superiority can be intuitively understood in the case of two clones that are consistently ranked the same way in all MCMC iterations, but are shifted by a random but shared offset.

**ctDNA detection.** We now describe how we formulate the task of ctDNA detection as Bayesian model selection. We compare two models. The first is a model hypothesizing tumour presence, $\mathbb{P}_1$, namely, the model described in Section 1.1. The second model, $\mathbb{P}_0$, is identical, except that the Dirichlet prior on $\rho$ is replaced by a point mass distribution on the normal population,

$$\rho \equiv (0, 0, \ldots, 0, 1).$$

In other words, $\mathbb{P}_0$ hypothesizes that no cancer cells are shedding detectable cfDNA.

Given a cfDNA sample $\mathcal{D} := (B_t, L_t)_{t=1}^{T}$, define the (base-10) log Bayes factor as:

$$B = \log_{10}\frac{\mathbb{P}_1(\mathcal{D})}{\mathbb{P}_0(\mathcal{D})}.$$

Then to perform model comparison we use the following thresholds to determine the ctDNA state of a cfDNA sample

$$f(B) = \begin{cases} \text{Detected} & B \geq 3 \\ \text{Inconclusive} & -3 < B < 3 \\ \text{Not detected} & B \leq -3. \end{cases}$$

**ctDNA genomic bin outlier probability.** Recall $z^{\text{rdr}} = (z_1^{\text{rdr}}, \ldots, z_T^{\text{rdr}})$ where $z_t^{\text{rdr}}$ denotes the RDR outlier indicator variable for genomic bin $t$ and $L_t = l_t$ denotes an observed RDR at genomic bin $t$. As stated previously, $z^{\text{rdr}}$ is analytically marginalized out during MCMC sampling therefore we must use the existing samples to obtain a Monte Carlo estimate of the postior mean of $z^{rdr}$

$$\hat{z}_t^{\text{rdr}} = \frac{1}{M}\sum_{m=1}^{M} \frac{\pi_{\text{rdr}}^{(m)} t_4(l_t|1,1)}{\pi_{\text{rdr}}^{(m)} t_4(l_t|1,1) + (1-\pi_{\text{rdr}}^{(m)}) t_{25}(l_t|\mu_t^{(m)}, \sigma^{(m)})}.$$

Similarly, we are able to use method to obtain outlier probabilities using the BAF data

$$\hat{z}_t^{\text{baf}} = \frac{1}{M} \sum_{m=1}^{M} \frac{\pi_{\text{baf}}^{(m)} f(b_t | d_t, 1, 1)}{\pi_{\text{baf}}^{(m)} f(b_t | d_t, 1, 1) + (1 - \pi_{\text{baf}}^{(m)}) f(b_t | d_t, \frac{p_t^{(m)}}{\gamma^{(m)}}, \frac{1 - p_t^{(m)}}{\gamma^{(m)}})}$$

where $f(b|d, \alpha, \beta)$ denotes the probability density function for a Beta Binomial distribution with mean $\frac{d\alpha}{\alpha+\beta}$ and variance $\frac{d\alpha\beta(\alpha+\beta+d)}{(\alpha+\beta)^2(\alpha+\beta+1)}$.

#### 2.2 Approximate cfClone posterior inference

We describe here the inference procedure used to approximate the posterior distribution of cfClone. Our inference methods overcome two challenges. First, we need not only samples from the posterior, but also robust approximation of marginal likelihoods $\mathbb{P}_i(\mathcal{D})$, $i \in \{0, 1\}$ needed for ctDNA detection via model selection. Second, we found empirically that cfClone posterior distributions can be *ill-conditioned* (i.e., some directions are highly constrained compared to other directions, causing inefficient simulation under naive schemes).

To address the first challenge, we use non-reversible parallel tempering[7] (NRPT) combined with the stepping stone estimator[8], implemented in the `Pigeons` software suite[9]. The model is implemented in the `Stan` Bayesian modelling language[10], and interfaced with `Pigeons` using the `BridgeStan` interface[11]. To address the second challenge, we first compute a Laplace approximation, and use the covariance of the Laplace approximation as the inverse mass matrix of the HMC NUTS algorithm (Hamiltonian Monte Carlo No-U-Turn Sampler)[12] used as the local exploration kernel of NRPT. This acts as a full rank preconditioner. The HMC NUTS implementation is provided by the `AdvancedHMC` package[13]. We also use the same Laplace approximation to initialize the NRPT algorithm and to provide an informed reference distribution.

Empirically, the cfClone posterior distribution appears to be log-concave for most post warm-up samples, however, this is not the case during the first iterations of the optimization problem required to compute the Laplace approximation. To address this issue, we used a regularized-Hessian version of Newton's method. Specifically, for iterates where the minimum eigenvalue of the negative Hessian was smaller than $10^{-5}$, we added to the Hessian a scaled identity matrix to ensure the Newton preconditioner is positive definite (identity matrix was scaled by two times the minimum eigenvalue plus $10^{-5}$). We used a straightforward extension of AutoGD[14], adding our regularized Newton preconditioner to vanilla AutoGD, to perform step size adaptation. This yields a robust optimization algorithm that does not require user tuning.

### 3 Supplementary Note: Semi-synthetic data simulation methods

In the following, we describe two methods to generate semi-synthetic cfDNA data in order to evaluate cfClone's ability to perform tumour fraction and clonal prevalence estimation and ctDNA detection. First we describe how to generate semi-synthetic data informed by scWGS using the infinite pool model in Section 3.1 followed by generating semi-synthetic data by mixing ctDNA bams in Section 3.3. The first generation method provides a setting in which ground truth tumour fraction and clonal prevelance are known. The second method allows us to evaluate cfClone on real patient derived cfDNA data to ensure that our model performs well in the cfDNA data setting.

#### 3.1 Simulating cfDNA using scWGS and a matched normal WGS

True tumour fraction and clonal prevalence of cfDNA samples are unknown. Therefore to systematically evaluate cfClone, we develop a simulator called the infinite pool model that uses scWGS and a matched normal WGS sample to simulate cfDNA. Our simulator requires binned read counts and clone assignments of cancer cells and a matched normal WGS sample. In brief, the model create in-silico mixtures of cell populations at fixed tumour fraction and clonal prevalences specified by $\rho$. That is, in contrast to Section 1, $\rho$ is not a reconstructed quantity but rather an input. The outputs of our simulator are total and haplotype binned read counts.

##### 3.1.1 Sampling binned read counts

Given a desired coverage $H \in \mathbb{R}^+$ the number of reads $N \in \mathbb{N}^+$ is computed by

$$N = \left\lceil \frac{\mathcal{G}}{\mathcal{R}} H \right\rceil \tag{4}$$

where $\mathcal{R} \in \mathbb{N}^+$ and $\mathcal{G} \in \mathbb{N}^+$ are constants denoting read length and human genome length given in base pairs (bp). Compute the ploidy $m_k \in \mathbb{R}$ for each cell population $k$ by

$$m_k = \frac{1}{T} \sum_t C_{tk}$$

We compute the number of reads from each cancer cell population $k$ as follows:

$$N_k = \left\lceil N \cdot \frac{\rho_k m_k}{\sum_{k'} \rho_{k'} m_{k'}} \right\rceil \tag{5}$$

Let $\bar{D}_i$ denote the normalized binned read counts of cell $i$ such that $\bar{D}_{it} = \frac{D_{it}}{\sum_{t'} D_{it'}}$. Let $Z_k$ denote the total number of cells assigned to population $k$ and $i_k$ denote a cell assigned to population $k$. In what follows, we will treat the inputted normal sample as a normal cell so that $Z_{K+1} = 1$ and "$\sum_{i_k}$" denotes the summation of cells assigned to cell population $k$.

$$M_k \sim \text{Multinomial}\big(N_k, \underbrace{(\frac{1}{Z_k}, \ldots, \frac{1}{Z_k})}_{Z_k \text{ cells}}\big) \tag{6}$$

$$P_k^{i_k} \sim \text{Dirichlet}(\bar{D}_{i_k}) \tag{7}$$

$$R_k^{i_k} | M_k^{i_k}, P_k^{i_k} \sim \text{Multinomial}(M_k^{i_k}, P_k^{i_k}) \tag{8}$$

$$R_k | \{R_k^{i_k}\}_{i_k} \sim \mathbb{1}\left[R_k | R_{kt} = \sum_{i_k} R_{kt}^{i_k}\right] \tag{9}$$

$$G | \{R_k\}_k \sim \mathbb{1}\left[G | G_t = \sum_k R_{kt}, \forall t \in [T]\right] \tag{10}$$

Equation 6 samples number of reads generated from $Z_k$ cells uniformly at random. Equation 7 samples a simplex $P_k^{i_k}$ that represents the probability of a read mapping to genomic bin $t$ informed by cell $i_k$. Equation 8 samples binned read counts generated from cell $i_k$. Equation 9 aggregates binned read counts for all cells in population $k$. Similarly, Equation 10 aggregates binned read counts across all cell populations $k$.

#### 3.2 Sampling haplotype binned read counts

Let $s_t \in [0, 1]$ denote the heterozygous SNP density of bin $t$ computed by the ratio between number of heterozygous SNPs and length of bin $t$ using the matched normal sample.

$$r_t^{i_k} \sim \mathbb{1}\left[r_t^{i_k} = \mathcal{R} \cdot s_t \cdot D_{i_k t}\right] \tag{11}$$

$$V_t^{i_k} \sim \text{Poisson}(r_t^{i_k}) \tag{12}$$

$$B_t^{i_k} \sim \text{Binomial}\left(V_t^{i_k}, \frac{C_{i_k t}^b}{C_{i_k t}}\right) \tag{13}$$

$$B_{kt} | \{B_t^{i_k}\}_{i_k} \sim \mathbb{1}\left[B_{kt} | B_{kt} = \sum_{i_k} B_t^{i_k}\right] \tag{14}$$

$$Q_t | \{B_{kt}\}_k \sim \mathbb{1}\left[Q_t | Q_t = \sum_k B_{kt}\right] \tag{15}$$

Equation 11 computes the expected number of reads to cover a heterozygous SNPs in genomic bin $t$ which will be proportional to read length $\mathcal{R}$ and SNP density $s_t$. Intuitively, this means that as the read length or SNP density increases then the probability of a read overlapping a heterzygous SNP will also increase. Equation 12 which samples the number reads that cover a heterzygous SNPs in bin $t$. Equation 13 then samples the number of reads matching the B allele in bin $t$. Note the ratio between the B haplotype and total copy numbers which is then used as the probability that a read will map to haplotype B and when $k = K + 1$ is the normal cell population is 0.5. Equation 14 sums haplotype counts across cell assigned to cell population $k$. Similarly, Equation 15, sums haplotype counts across cell populations.

##### 3.2.1 Simulator validation

We first validated the infinite pool model by verifying that the simulated cfDNA samples are concordant with the observed cfDNA data for HGSOC patient CID12347. We performed inference with cfClone to obtain tumour fraction and clonal prevalence estimates for each cfDNA sample then used these estimates as inputs for the infinite pool model. We simulated 100 replicates to verify the observed data was contained in the 95% quantile interval. These results are shown in Figure 3. 30% of genomic bins are contained outside the 99% quantile intervals for the RDR data and 1% of genomic bins are contained outside the 99% qunatiles interval for the BAF data. The largest source of deviation is chromosome 19 where 68% of genomic bins are not contained however we hypothesize this is due to ctDNA specific data biases that are not captured in scWGS. Of the genomic bins that are outside the 99% quantile, the average distance from the closest bounds is 0.018 with a standard deviation of 0.020 where the max is 0.210 and occurs on Chromosome 19.

#### 3.3 In-silico mixing of patient derived ctDNA samples

Our goal is to create a consistency check on tumour fraction estimation outputted by cfClone on patient derived ctDNA samples to ensure that cfClone is robust to ctDNA specific data artifacts. We do so by mixing two sets of reads that we call "end points" index by $\beta \in \{0, 1\}$. We mix reads from the two end points at proportions denoted by $\beta \in (0, 1)$ to generate intermediate mixtures as described in Section 3.3.1. Then, using a theoretical framework described in Section 3.3.2, we show how to use $\rho^0$, $\rho^1$, and $m_k$, cfClone's prevalence estimates obtained on the endpoints and clone copy number profiles, to validate inferred $\rho^\beta$ on the intermediate mixture.

##### 3.3.1 Generating mixtures of end point reads

We aim to generate a mixture of end point reads at proportion $\beta \in (0, 1)$ with coverage $H^\beta$. We first compute the total number of reads $N^\beta$ required to obtain a coverage $H^\beta$ using Equation 4 as in Section 3.1.1. Next we compute the number of reads from each end point that will make up the intermediate mixture

$$N^0 = \lceil N^\beta(1-\beta) \rceil$$
$$N^1 = \lceil N^\beta \beta \rceil .$$

Using *samtools view –subsample* we subsample each end point to $N^0$ and $N^1$ reads using arguments $\frac{N^0}{R^0}$ and $\frac{N^1}{R^1}$ where $R^0$ and $R^1$ denote the total number of pair end reads available in each end point after filtering for quality metrics using *samtools view*. Finally, we merge the down sampled set of reads using *samtools merge*.

##### 3.3.2 Predicting cell population prevalences

Using inputs $\rho^0$, $\rho^1$, and $m_k$ we derive a theoretical framework that allows us to estimate $\rho^\beta$. First, we define $r$ which is a simplex of length $K + 1$ that denotes the read proportion of each cancer cell population and normal population. Formally, $r = (r_1, \ldots, r_K, r_{K+1})$ denotes a simplex-valued parameter of size $K + 1$, $\sum_{k=1}^{K+1} r_k = 1$, where the last component, $n = K + 1$, is reserved for the normal cell prevalence. One can think of $r$ as being the "read space" equal to the "cell space" $\rho$. The way we mix reads in Section 3.3.1 satisfies the following equation we refer to as the "mixing" equation:

$$r_k^\beta = \beta r_k^1 + (1-\beta) r_k^0. \tag{16}$$

Equation 5 in Section 3.1.1 is equivalent to

$$\frac{r_k^\beta}{r_{k'}^\beta} = \frac{m_k}{m_{k'}} \frac{\rho_k^\beta}{\rho_{k'}^\beta} \quad \forall k, k' \text{ s.t } k \neq k. \tag{17}$$

which we will refer to as the "shedding" equation. For both $\rho$ and $r$ we will use the shedding Equation 17 with $k' = n$ to define their "ratio-to-normal". More formally, for a vector $\tilde{x} = (\tilde{x}_1, \ldots, \tilde{x}_n) \in \mathbb{R}^{K+1}$ define $\tilde{x}_{\cdot:n} = (\tilde{x}_{1:n}, \ldots, \tilde{x}_{n-1:n})$ where $\tilde{x}_{k:n} = \tilde{x}_k/\tilde{x}_n$. Notice, there is a one-to-one mapping between $\tilde{x}_{\cdot:n}$ and

$$x = (\frac{\tilde{x}_1}{\sum_k \tilde{x}_k}, \ldots, \frac{\tilde{x}_n}{\sum_k \tilde{x}_k}).$$

The forward map follows by definition of $\tilde{x}_{\cdot:n}$ and the reverse map is given by $x_k = x_n \tilde{x}_{k:n}$ where

$$x_n = \left(1 + \sum_{k=1}^{n-1} \tilde{x}_{k:n}\right)^{-1}.$$

Using the mixing Equation 16, shedding Equation 17 and ratio-to-normal of $\rho$ and $r$ we can derive our prediction method into five steps. First for $\beta \in \{0, 1\}$ transform $\rho^\beta$ into $r^\beta$ use the shedding Equation 17. Next for intermediate $\beta \in (0, 1)$ use the mixing Equation 16 to compute $r^\beta$. Thirdly, transform $r^\beta$ to $\tilde{r}^\beta_{\cdot:n}$ by definition, followed by using the shedding Equation 17 to transform $\tilde{r}^\beta_{\cdot:n}$ to $\tilde{\rho}^\beta_{\cdot:n}$. Finally, use the reverse mapping to transform $\tilde{\rho}^\beta_{\cdot:n}$ to $\rho^\beta$. Ultimately, Proposition 2 shows that we can decompose the prediction equation into a linear mixing and correction term. This equation can be interpreted as the correction required when if one naively predicts mixture prevalences linearly without accounting for the ploidy of cell populations.

**Proposition 2.** *The prediction equation can be expressed as follows:*

$$\rho_k^\beta = \underbrace{\left[\beta \rho_k^1 + (1-\beta)\rho_k^0\right]}_{\textit{naive linear prediction}} + \underbrace{(\rho_k^1 - \rho_k^0)\beta(1-\beta)\frac{(\mathbb{m}^0 - \mathbb{m}^1)}{\beta \mathbb{m}^0 + (1-\beta)\mathbb{m}}}_{\textit{ploidy correction}} \tag{18}$$

*where* $\mathbb{m}^0 = \sum_k m_k \rho_k^0$ *and* $\mathbb{m}^1 = \sum_k m_k \rho_k^1$.

*Proof.* Using the shedding Equation 17, we obtain the following

$$\rho_k^\beta = \frac{r_k^\beta / m_k}{\sum_{k'} (r_{k'}^\beta / m_{k'})} \quad \text{for} \quad k \in [K]. \tag{19}$$

which we will now derive expressions for the numerator and denominator and obtained the decomposed form in Proposition 2. Let $\mathbb{m}^0 = \sum_k m_k \rho_k^0$ and $\mathbb{m}^1 = \sum_k m_k \rho_k^1$ so that $r_k^0 = m_k \rho_k^0 / \mathbb{m}^0$ and $r_k^1 = m_1 \rho_k^1 / \mathbb{m}^1$. Following Steps 1 and 2 above we obtain the following expressions for the numerator and denominator:

$$\frac{r_k^\beta}{m_k} = \beta \frac{\rho_k^1}{\mathbb{m}^1} + (1-\beta)\frac{\rho_k^0}{\mathbb{m}^0} \quad \text{for} \quad k \in [K].$$

$$\sum_{k'} \frac{r_{k'}}{m_{k'}} = \beta \frac{\sum_{k'} \rho_{k'}^1}{\mathbb{m}^1} + (1-\beta)\frac{\sum_{k'} \rho_{k'}^0}{\mathbb{m}^1} = \frac{\beta}{\mathbb{m}^1} + \frac{(1-\beta)}{\mathbb{m}^0}.$$

We now combine the expression for the numerator and the denominator

$$\rho_k^\beta = \frac{\beta \frac{\rho_k^1}{\mathbb{m}^1} + (1-\beta)\frac{\rho_k^0}{\mathbb{m}^0}}{\frac{\beta}{\mathbb{m}^1} + \frac{(1-\beta)}{\mathbb{m}^0}} = \frac{\beta \rho_k^1 \mathbb{m}^0 + (1-\beta)\rho_k^0 \mathbb{m}^1}{\beta \mathbb{m}^0 + (1-\beta)\mathbb{m}^1}.$$

Now subtract $A = \beta\,\rho_k^1 + (1-\beta)\,\rho_k^0$ then divide by $D = \beta \mathbb{m}^0 + (1-\beta)\mathbb{m}^1$

$$\rho_k^\beta - A = \frac{\beta\,\rho_k^1 \mathbb{m}^0 + (1-\beta)\,\rho_k^0 \mathbb{m}^1 - AD}{D} \tag{20}$$

Expanding $AD$ and using $\beta - \beta^2 = \beta(1-\beta) = (1-\beta) - (1-\beta)^2$ the numerator reduces to

$$\beta(1-\beta)\,\rho_k^1\big(\mathbb{m}^0 - \mathbb{m}^1\big) - \beta(1-\beta)\,\rho_k^0\big(\mathbb{m}^0 - \mathbb{m}^1\big) = \beta(1-\beta)\big(\rho_k^1 - \rho_k^0\big)\big(\mathbb{m}^0 - \mathbb{m}^1\big). \tag{21}$$

Hence

$$\rho_k^\beta - A = \frac{\beta(1-\beta)(\rho_k^1 - \rho_k^0)(\mathbb{m}^0 - \mathbb{m}^1)}{D} \tag{22}$$

which gives the claimed decomposition. □

### 4 Supplementary Note: Synthetic results

In the following experiment we run cfClone for 12 rounds and 5 chains leveraging the Laplacian approximation to precondition the HMC kernels and as the reference distribution. The clone copy number profiles are inputted into cfClone for any clone with positive prevalence. We run ichorCNA using the code from their publication[15]. For each generated dataset, we output a wig file of the binned read counts to pass into ichorCNA so that their preprocessing procedure that includes GC, mappability, and panel of normal (PoN) correction are performed. We set the binsize of ichorCNA to be the same as cfClone at $500K$kb. Following ichorCNA's suggestion on their paper and Github we use two different initialization settings depending the on the dataset generation settings. For the standard scheme we use following purity $n \in \{0.35, 0.45, 0.5, 0.6, 0.7, 0.8, 0.95, 0.99\}$ and ploidy $\phi \in \{2, 3, 4\}$ values to initialize their EM algorithm. For the low ctDNA scheme we use following purity $n \in \{0.95, 0.99, 0.995, 0.999\}$, ploidy $\phi \in \{2\}$ and set a max copy number of 2 and exclude subclonal events.

#### 4.1 Benchmarking cfClone tumour fraction estimation against ichorCNA using of three different cancer types: FL, DLBCL, and HGSOC

We evaluate tumour fraction estimation using in-silico data where true tumour fraction and clonal prevalences are known. Clone copy number profiles are inputted into cfClone for any clones with positive prevalence so that cfClone has full information on tumour heterogeneity. For each cancer type we place uniform clonal prevalence on existing clones. We generate data using 16 different tumour fraction values between 0% to 75% and 6 different coverage settings between $0.1X$ and $100X$ to understand the affect of sequencing depth on estimation. We repeat this process using scWGS of three different cancer types in order of increasing copy number instability: FL, DLBCL, and HGSOC.

We use the following metrics to evaluate and benchmark cfClone tumour content estimation. Let $t \in T$ denote the set of expected tumour contents and $\hat{t}$ denote cfClone mean tumour content estimate. The mean and max absolute error is computed by $\text{MAE} = \sum_{t\in T} |t - \hat{t}|$ and $\text{MaxAE} = \max_{t\in T} |t - \hat{t}|$, respectively. Notice that the mean and max are over the set of all tumour fraction values so that we may summarize the results per level of coverage. The relative absolute error is computed by $r = \frac{|t-\hat{t}|}{t}$.

For each cancer type, at coverage $0.1X$ we observe the largest mean absolute error for tumour fraction estimation of 0.8%, 0.5%, and 0.7% for FL, DLBCL, and HGSOC, respectively. As coverage increases from $0.1X$ to $100X$ the mean absolute error decreases to 0.2%, 0.2%, and 0.1% for FL, DLBCL, and HGSOC, respectively (Supplemental Figures 4- 5, 6- 7, and 8- 9). We repeated the above analysis for 10 data replicates and observed the largest mean absolute errors again occur at $0.1X$ with values of 0.8%, 0.4% and 0.6% for FL, DLBCL, and HGSOC. Increasing coverage from $0.1X$ to $100X$ saw a decrease in the average of the mean absolute error across data replicates to 0.2%, 0.2% and 0.1% for FL, DLBCL, and HGSOC (Supplemental Figures 5, 7 9).

Table 2 shows cfClone outperforms ichorCNA as measured by mean and max absolute error for all cancer types at coverage $100X$ however similar results hold for the other coverage settings. cfClone performed similarly on HGSOC and DLBCL achieving mean absolute errors of 0.1% and 0.2%, respectively. In comparison, ichorCNA achieved a mean absolute error of 1.7% on HGSOC and 2.4% on DLBCL, both of which are an order of magnitude larger than cfClone. In general ichorCNA had large max absolute errors of 3%, 22%, and 5% on FL, DLBCL, and HGSOC, respectively. cfClone and ichorCNA performed most similarly on FL achieving mean absolute errors of 0.2% and 0.3%, respectively.

#### 4.2 cfClone provides sensitive ctDNA detection on in-silico data generated from HGSOC, DLBCL, and FL

Using the same data generating process as described in Section 4.1. Next we evaluated cfClone's ability to detect ctDNA using data generated with the infinite pool model using scWGS data from FL, DLBCL and HGSOC cancer patients. We vary coverage between $0.1X$ and $100X$ and tumour content between 0 and 75%. For each unique pair of coverage and tumour content we generate 10 cfDNA data replicates, then perform ctDNA detection as described in Section 2.1. Overall, as the coverage increases cfClone becomes more sensitive to ctDNA detections for each cancer type (Supplemental Figures 10 11 12). For DLBCL and HGSOC, with high sequencing depth ($100X$), cfClone was able to detect ctDNA for 100% of generated datasets at a tumour content of 0.1%. Conversely, at a low sequencing depth ($0.1X$), cfClone was able to detect ctDNA for 100% of generated datasets when tumour content was at least 2%, for DLBCL and HGSOC. As a control we simulated a pure normal sample and observed cfClone had a 0% false detection rate for FL, DLBCL, and HGSOC across all coverages evaluated (treating inconclusive outcome as a null result).

#### 4.3 Tumour fraction and clonal prevalence estimation on in-silico mixture of FL and DLBCL cancers

Next we evaluate cfClone clone prevalence accuracy on a mixture of FL and DLBCL cancers using data generated with the infinite pool model. We generate data using 16 different tumour fraction values between 0% to 75% and 6 different coverage settings between $0.1X$ and $100X$ to understand the affect of sequencing depth on estimation. We input three clones FL (A), DLBCL-00 (B), and DLBCL-01 (C) with fixed clonal prevalence of 0.5, 0.25, and 0.25, respectively. Results are shown in Figure 13 and the clone profiles are depicted in Figure 14.

At coverages $0.1X$ and $100X$ and tumour content $\geq$ 10% cfClone is able to capture the true clone prevalence within its 95% HDI intervals for subsequent tumour contents settings. At tumour content 0 and coverage $0.1X$ cfClone shows uncertainty regarding clone prevalence estimates with large overlapping HDI intervals. Additionally, at tumour content 0. and coverage $100X$ cfClone still shows uncertainty but prefers the clone that is most similar to the normal copy profile (Clone A). At coverages $0.1X$ and tumour contents $\leq$ 10% cfClone shows a large amount of uncertainty regarding the clone prevalence estimates and increasing coverage to $100X$ decreases uncertainty as shown by a decrease in the HDI interval widths. At the highest coverage of $100X$ and tumour content 0.25%, cfClone provides a confident on the tumour content estimate, but uncertain regarding clone prevalence estimates. This is intuitive, as it suggests that estimating tumour fraction is an easier problem than clone prevalence, with accurate estimates of the latter requiring high coverage in the low tumour fraction regime.

#### 4.4 cfClone provides consistent tumour fraction estimations on patient derived cfDNA and tissue samples

Next we evaluate cfClone tumour fraction estimation using mixtures of patient derived cfDNA and tissue samples. We describe our method to mix bam files in Section 3.3 and refer the reader to Section 3.3.2 for a derivation of the predicted tumour fraction. A total of four datasets are generated where two datasets are generated by mixing cfDNA samples from a single patient and the remaining two are generated by mixing tissue samples from a single patient. A total of three cfDNA samples are uniquely paired to generate two datasets of intermediate ctDNA mixtures. In particular, we mix a cfDNA sample estimated to contain near zero tumor content with two cfDNA samples estimated to contain tumor content. Similarly, three tissue samples are paired to generate two datasets of intermediate tissue mixtures. All intermediate mixtures are fixed to have a coverage of $30X$. Figure 15 shows the estimation of the tumor fraction for the four generated as a function of mixing proportion $\beta \in [0, 1]$. We observe that when mixing pairs of patient derived ctDNA and tissue samples that tumour content estimation is concordant with the theoretical prediction derived from our mixing framework. Subfigures **a** and **b** show results for mixtures of ctDNA data. The mean and max absolute errors are 0.065%, 0.088%, and 0.97%, 1.4%, respectively. We note that **b** seems to have a systematic bias to slightly overestimate tumor content. Subfigures **c** and **d** depict the results for mixtures of tissue samples. The mean and max absolute errors are 0.12%, 0.15%, and 1.6%, 2.5%, respectively.

### 5 Supplementary figures

7 Relative error of tumour content estimation as a function of expected tumour content for semi-synthetic data generated from scWGS of a DLBCL patient and their matched normal . 20
8 Tumour content estimation as a function of expected tumour content for semi-synthetic data generated from scWGS of a HGSOC patient and their matched normal . . . . . . . . . . . . . 21
9 Relative error of tumour content estimation as a function of expected tumour content for semi-synthetic data generated from scWGS of a HGSOC patient and their matched normal . 22
10 Proportion of data replicates where ctDNA was detected by cfClone for semi-synthetic data generated from scWGS of a FL patient and a matched normal . . . . . . . . . . . . . . . . . . 23
11 Proportion of data replicates where ctDNA was detected by cfClone for semi-synthetic data generated from scWGS of a DLBCL patient and a matched normal . . . . . . . . . . . . . . . 24
12 Proportion of data replicates where ctDNA was detected by cfClone for semi-synthetic data generated from scWGS of a HGSOC patient and a matched normal . . . . . . . . . . . . . . . 25
13 Tumour content and clone prevalence estimation on semi-synthetic data generated using scWGS data from FL and DLBCL cancers and a matched normal sample . . . . . . . . . . . 26
14 Clone total copy-number and BAF profiles for the FL and DLBCL clones obtained from scWGS 27
15 Inferred mean tumour fraction from cfClone and predicted tumour fraction by mathematical modelling for in-silico mixtures generated from cfDNA and bulk tumor BAMs . . . . . . . . . 28
16 Validation on the MSK-SPECTRUM dataset . . . . . . . . . . . . . . . . . . . . . . . . . 29

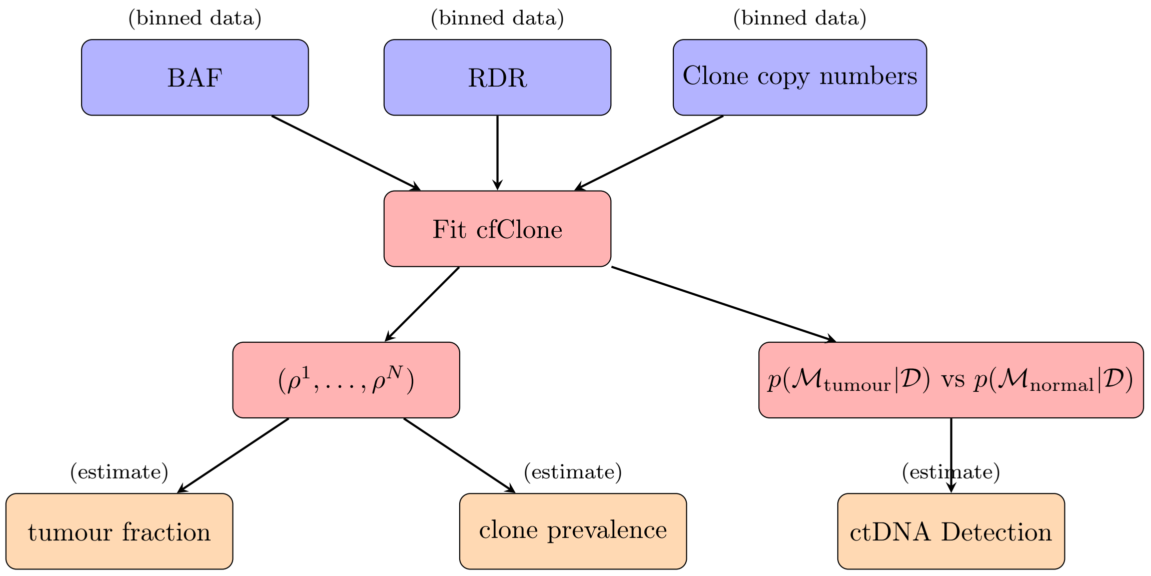

Figure 1: Overview of *cfClone* workflow used for benchmarking and evaluation on synthetic datasets. Purple nodes denote binned genomic data inputted into the model, red denotes model, inference, and parameters of interest, orange nodes denote key values used for analysis.

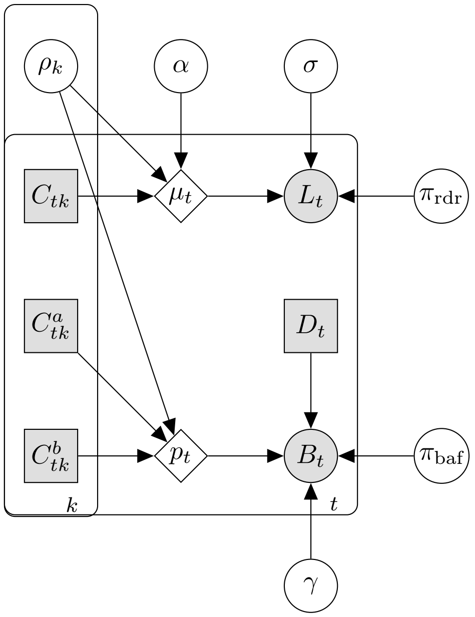

Figure 2: Probabilistic graphical model of cfClone. Shaded square nodes denote input data, shaded circle nodes denote observed data modeled with a likelihood, unshaded circle nodes denote latent variables, diamonds denote deterministic functions of their inputs shown as directed arrows. There are two plates $k \in [K+1]$ and $t \in [T]$ that denote the population index and genomic bins respectively.

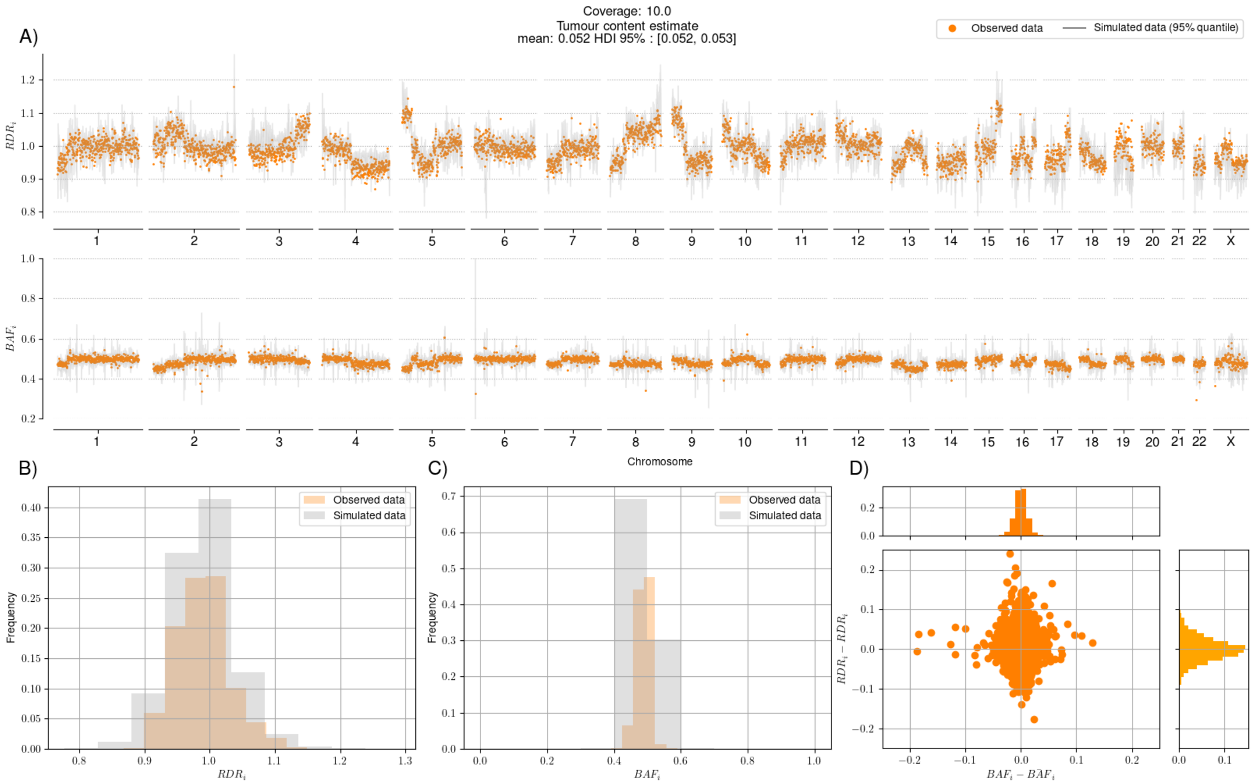

Figure 3: **A)** RDR and BAF for each genomic bin simulated using the infinite pool model. Orange denotes data from an observed HGSOC ctDNA sample and grey denotes the 99% quantile interval of samples generated via the infinite pool model for each genomic bin. **B) and C)** Historgrams of the observed ctDNA data (orange) and infinite pool samples (grey) for RDR and BAF data. **D)** Scatter plot of difference between observed data and mean of infinite pool samples for each genomic bin.

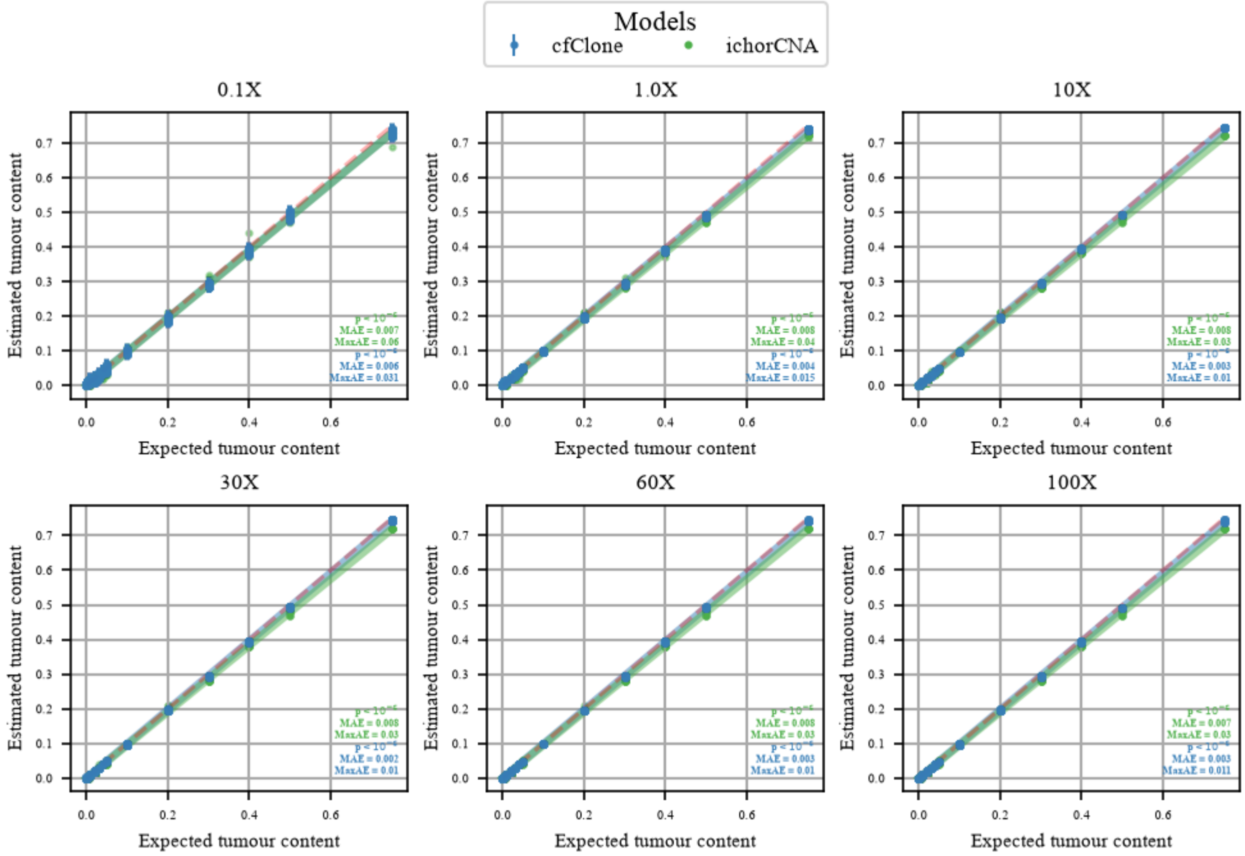

Figure 4: Tumour content estimation as a function of expected tumour content for data generated with the infinite pool model inputting scWGS of a FL patient and their matched normal. Coverages are varied for 6 values between $0.1X$ to $100X$ and tumour content is varied for 16 values between 0% to 75%. OLS regression was fit for cfClone using the mean estimated and expected tumour content. Circles indicate the mean tumour content estimate with whiskers showing the upper and lower bounds of the 95% HDI interval.

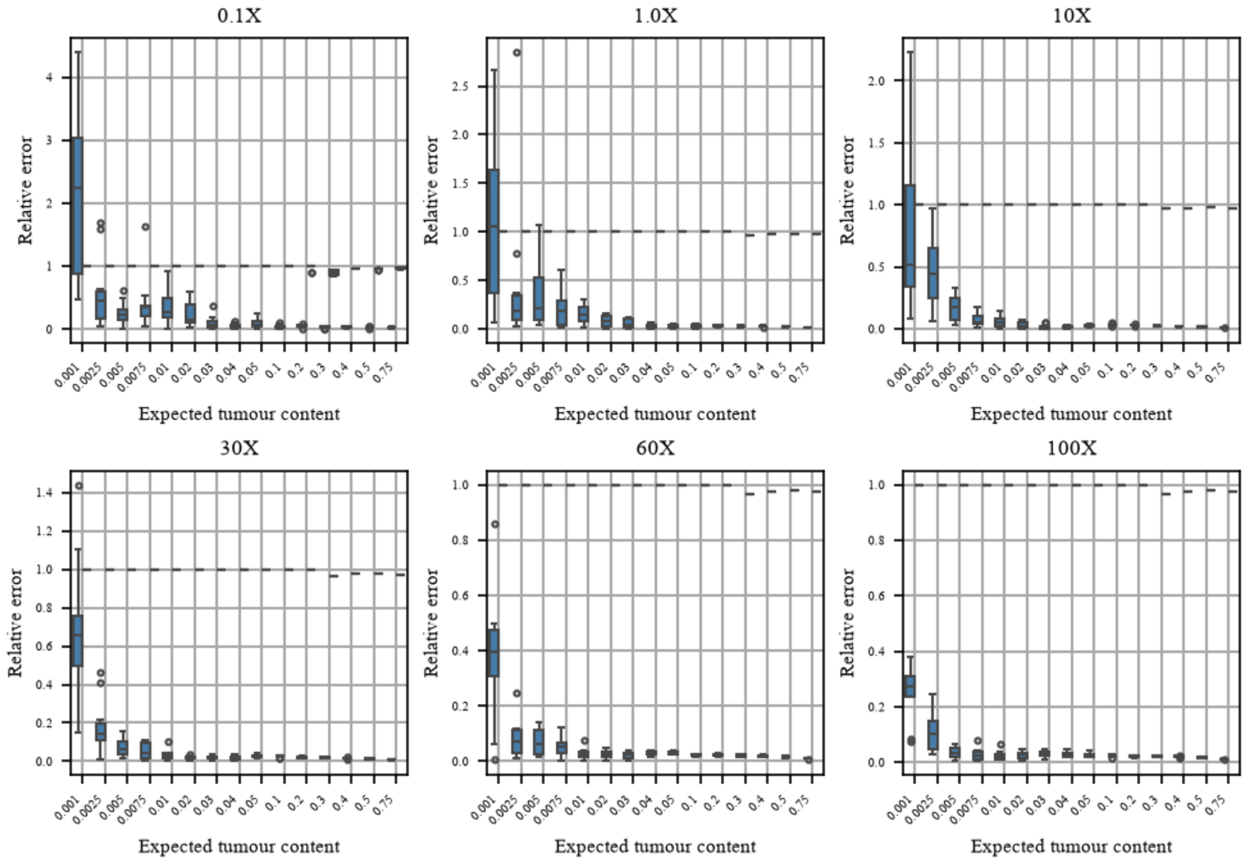

Figure 5: Relative error of tumour content estimation as a function of expected tumour content for each data replicate in log scale for data generated with the infinite pool model inputting scWGS of a FL patient and their matched normal. Coverages are varied for 6 values between $0.1X$ to $100X$ and tumour content is varied for 16 values between 0% to 75%. Error is measured by the difference between the mean tumour content estimate and the expected tumour content estimate for each of the 10 data replicates. Relative error is computed as the absolute difference between the mean and expected tumour content normalized by the expected tumour content. The results for cfClone are indicated in blue, while the results for ichorCNA are indicated in green.

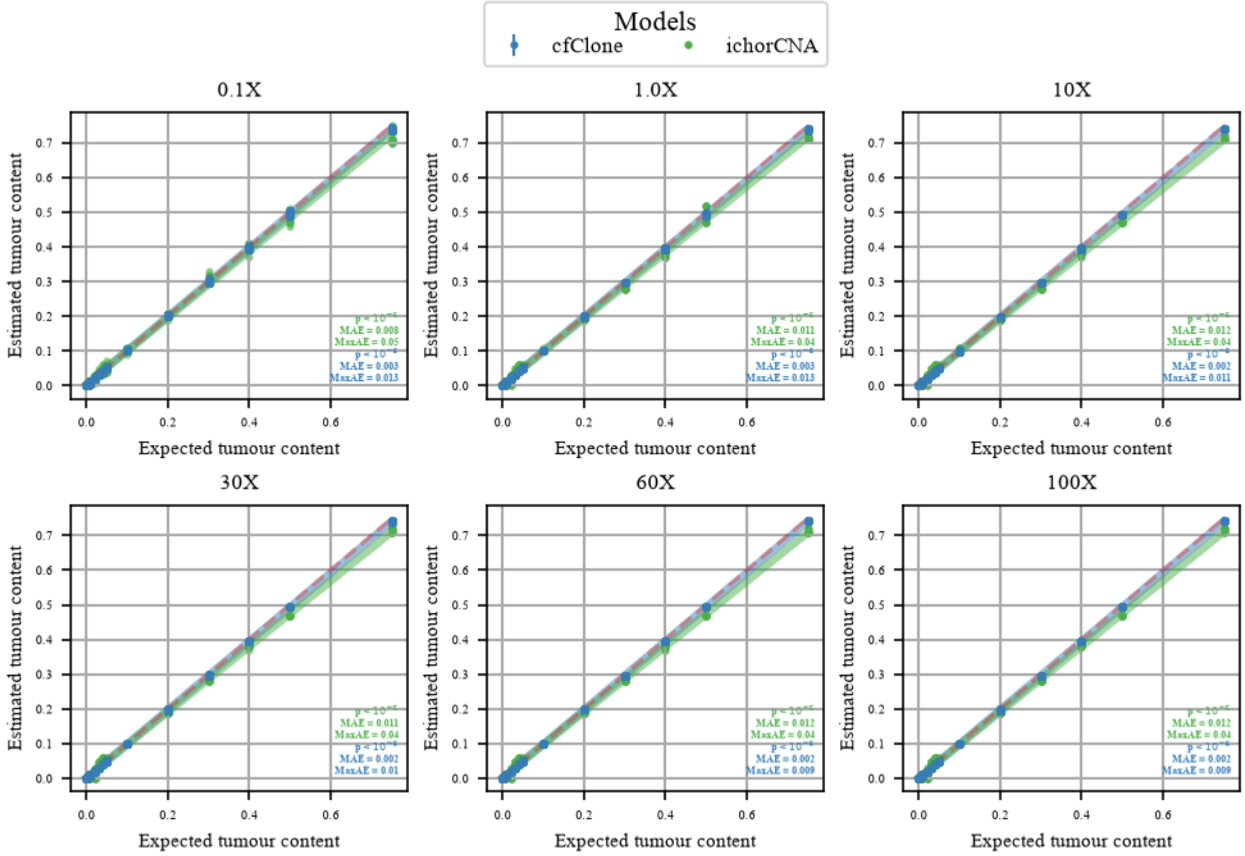

Figure 6: Tumour content estimation as a function of expected tumour content for data generated with the infinite pool model inputting scWGS of a DLBCL patient and their matched normal. Coverages are varied for 6 values between $0.1X$ to $100X$ and tumour content is varied for 16 values between 0% to 75%. OLS regression was fit for cfClone using the mean estimated and expected tumour content. Circles indicate the mean tumour content estimate with whiskers showing the upper and lower bounds of the 95% HDI interval.

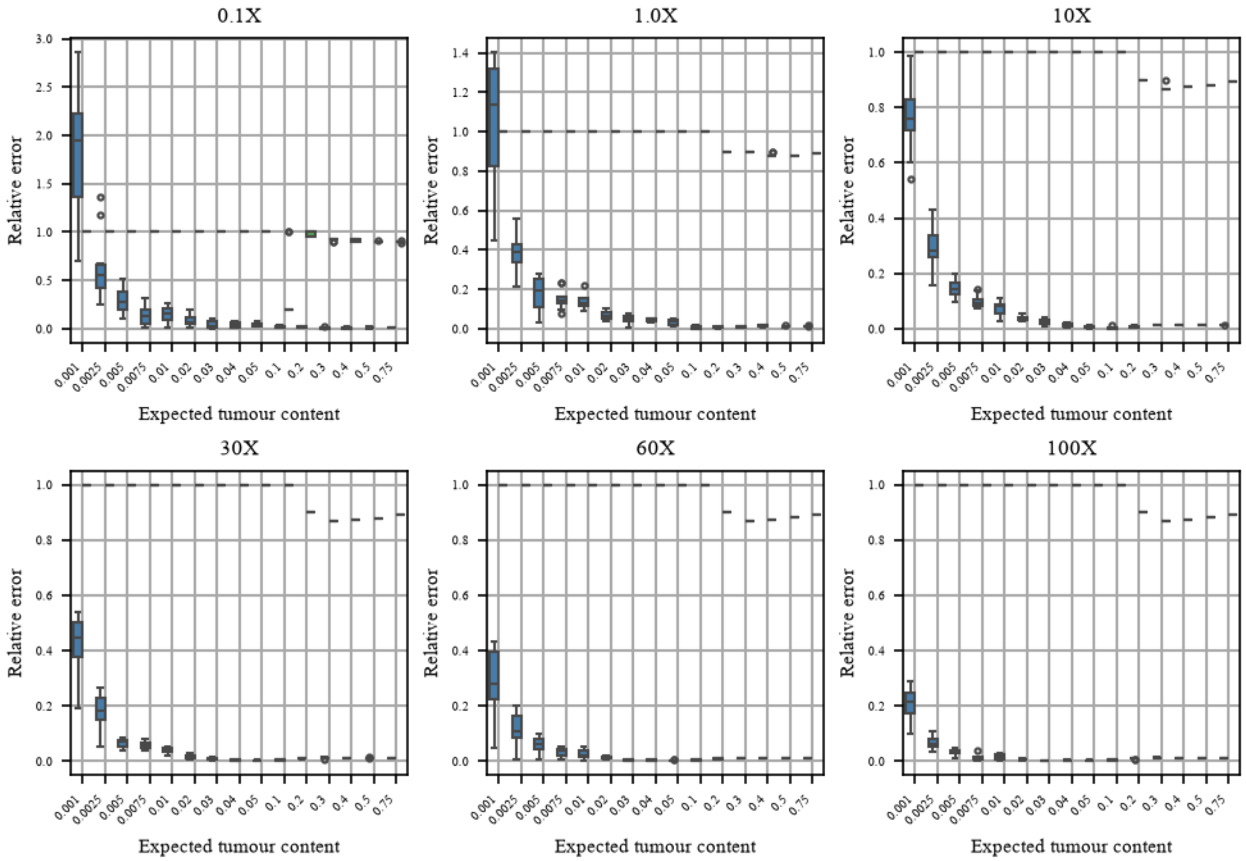

Figure 7: Relative error of tumour content estimation as a function of expected tumour content for each data replicate in log scale for data generated with the infinite pool model inputting scWGS of a DLBCL patient and their matched normal. Coverages are varied for 6 values between $0.1X$ to $100X$ and tumour content is varied for 16 values between 0% to 75%. Error is measured by the difference between the mean tumour content estimate and the expected tumour content estimate for each of the 10 data replicates. Relative error is computed as the absolute difference between the mean and expected tumour content normalized by the expected tumour content. The results for cfClone are indicated in blue, while the results for ichorCNA are indicated in green.

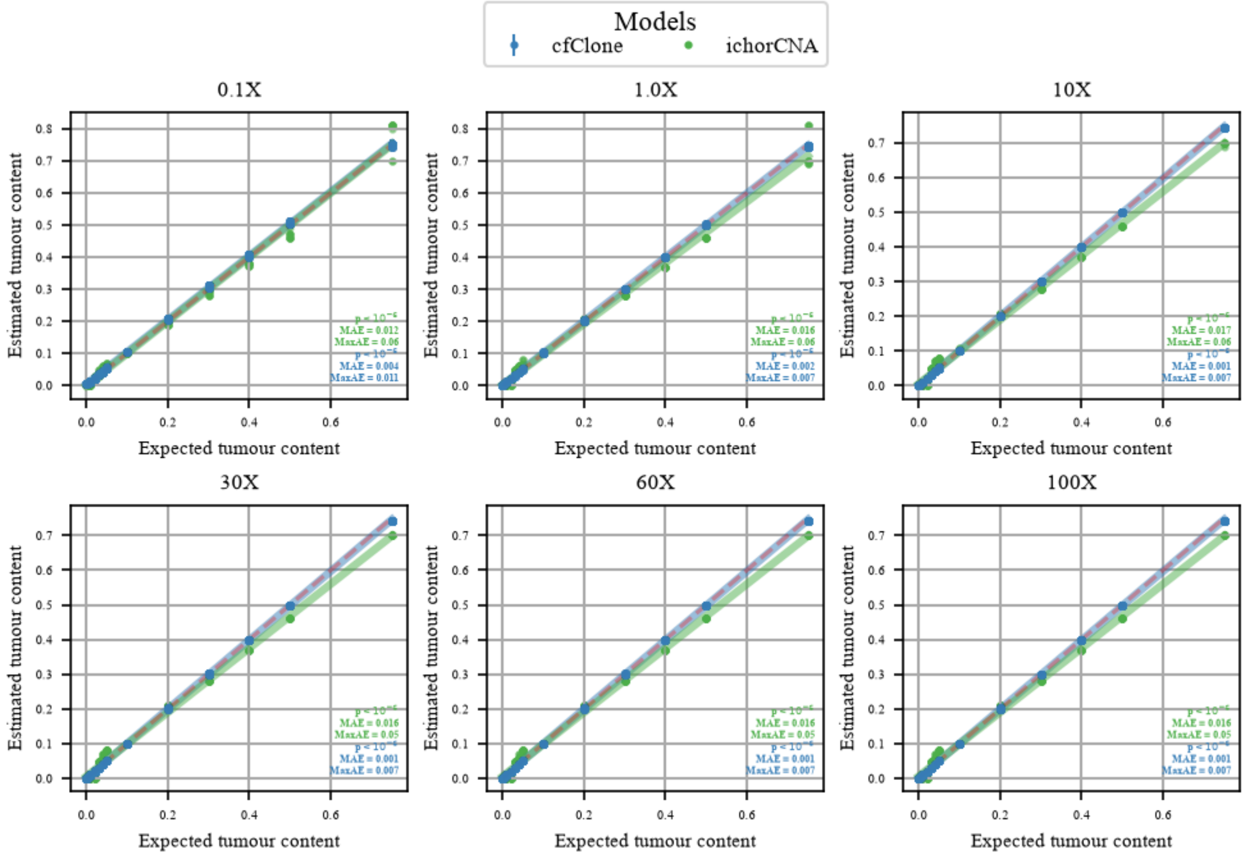

Figure 8: Tumour content estimation as a function of expected tumour content for data generated with the infinite pool model inputting scWGS of a HGSOC patient and their matched normal. Coverages are varied for 6 values between $0.1X$ to $100X$ and tumour content is varied for 16 values between 0% to 75%. OLS regression was fit for cfClone using the mean estimated and expected tumour content. Circles indicate the mean tumour content estimate with whiskers showing the upper and lower bounds of the 95% HDI interval.

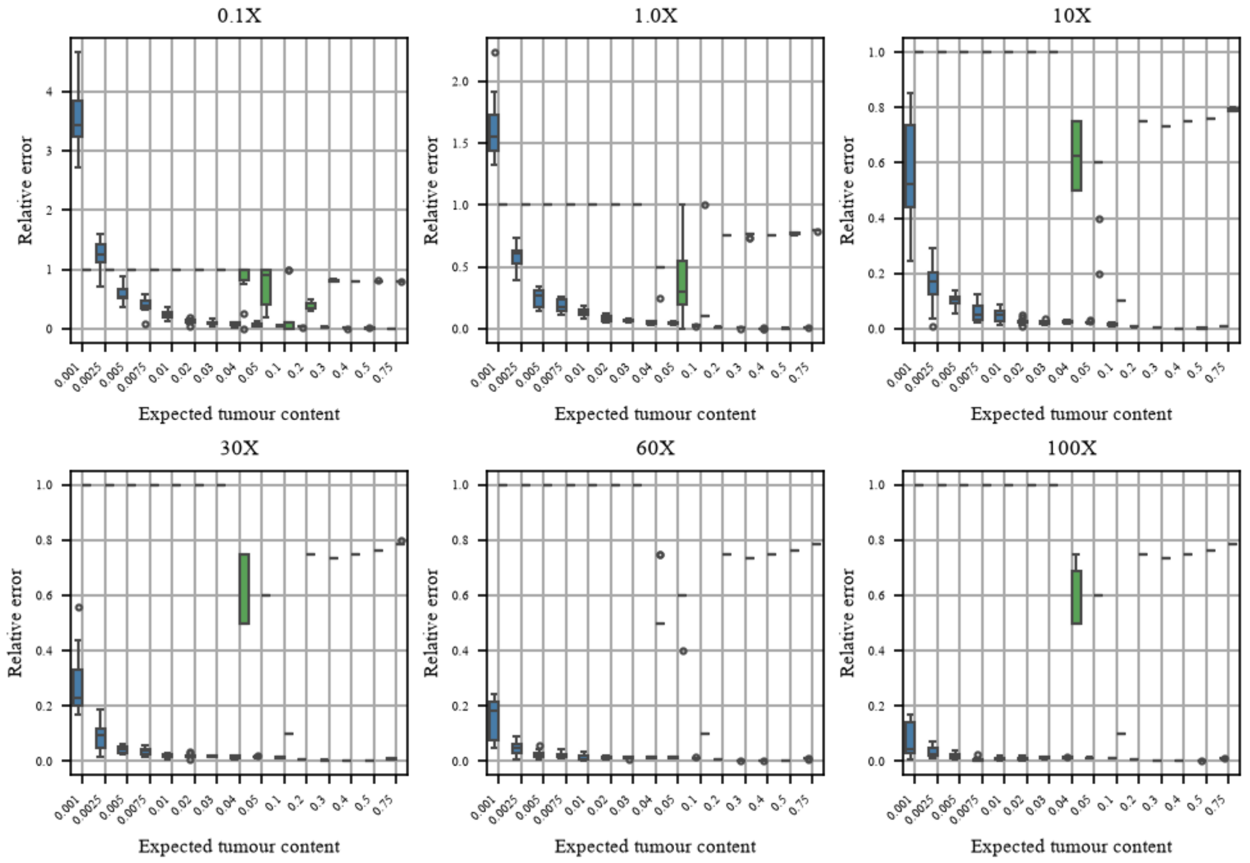

Figure 9: Relative error of tumour content estimation as a function of expected tumour content for each data replicate in log scale for data generated with the infinite pool model inputting scWGS of a HGSOC patient and their matched normal. Coverages are varied for 6 values between $0.1X$ to $100X$ and tumour content is varied for 16 values between 0% to 75%. Error is measured by the difference between the mean tumour content estimate and the expected tumour content estimate for each of the 10 data replicates. Relative error is computed as the absolute difference between the mean and expected tumour content normalized by the expected tumour content. The results for cfClone are indicated in blue, while the results for ichorCNA are indicated in green.

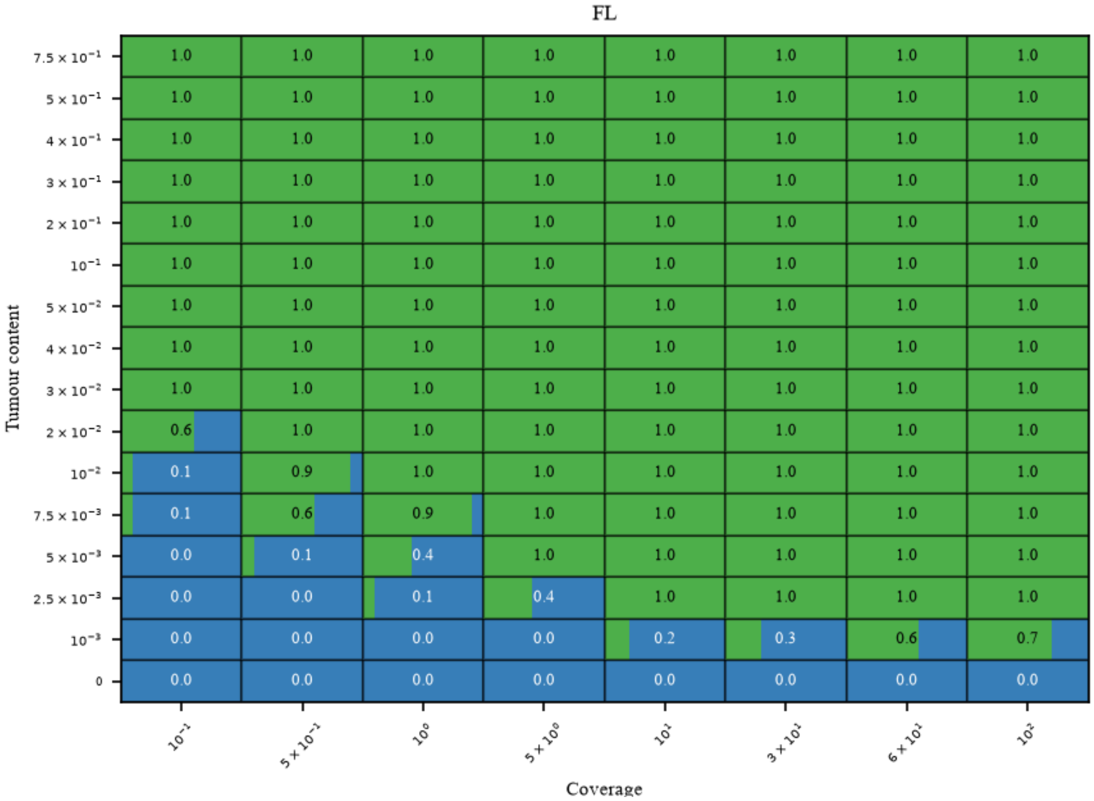

Figure 10: Proportion of data replicates where ctDNA was detected by cfClone when data was generated with the infinite pool model inputting scWGS of a FL patient and a matched normal. Coverages varied for 8 values between $0.1X$ and $100X$ and tumour content was varied for 16 values between 0% and 75%.

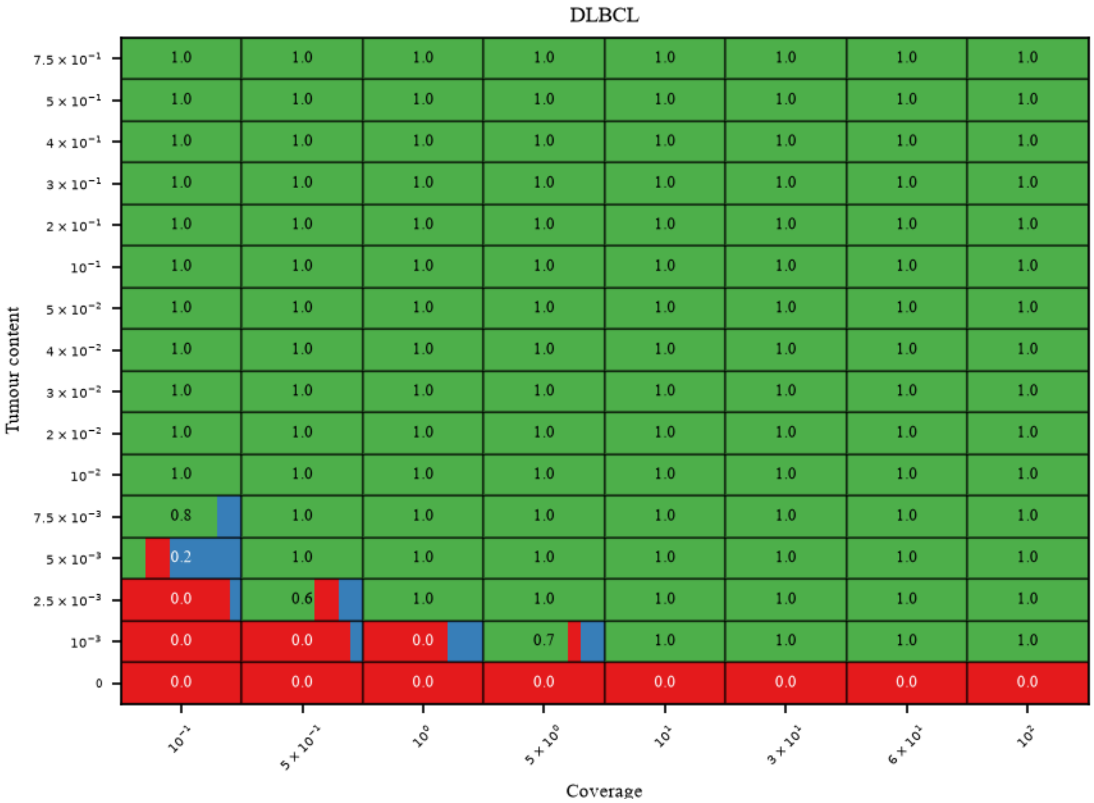

Figure 11: Proportion of data replicates where ctDNA was detected by cfClone when data was generated with the infinite pool model inputting scWGS of a DLBCL patient and a matched normal. Coverages varied for 8 values between $0.1X$ and $100X$ and tumour content was varied for 16 values between 0% and 75%.

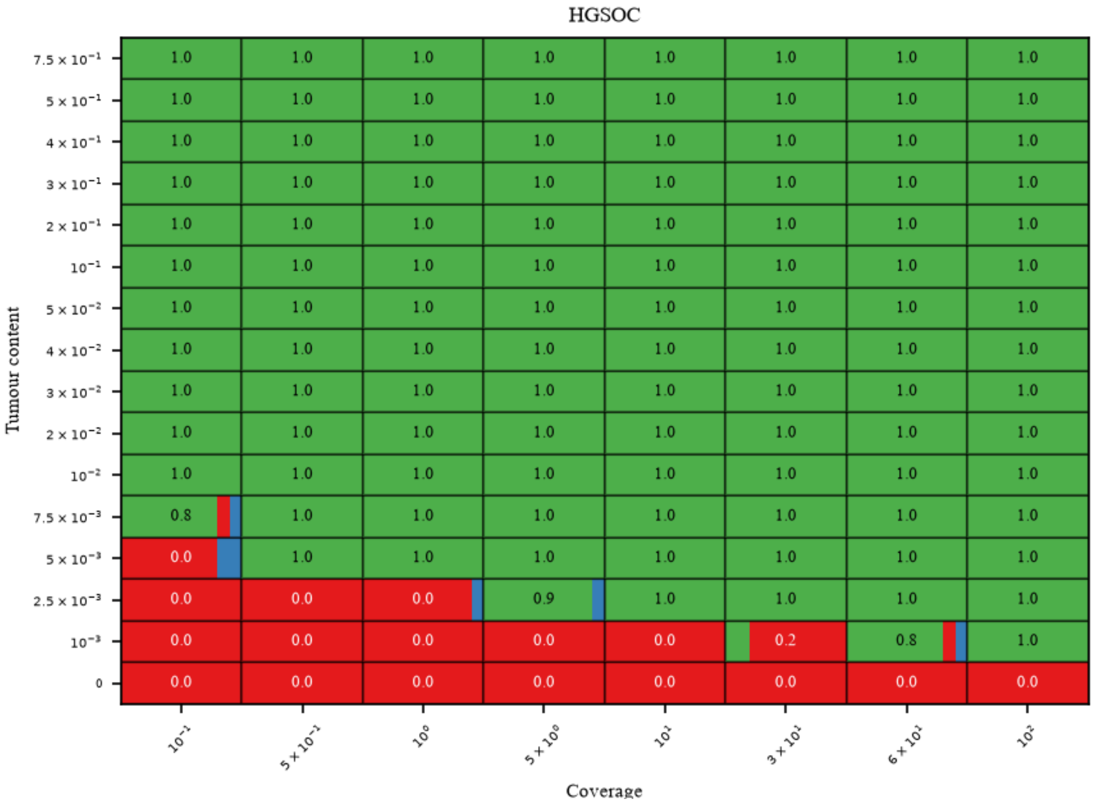

Figure 12: Proportion of data replicates where ctDNA was detected by cfClone when data was generated with the infinite pool model inputting scWGS of a HGSOC patient and a matched normal. Coverages varied for 8 values between $0.1X$ and $100X$ and tumour content was varied for 16 values between 0% and 75%.

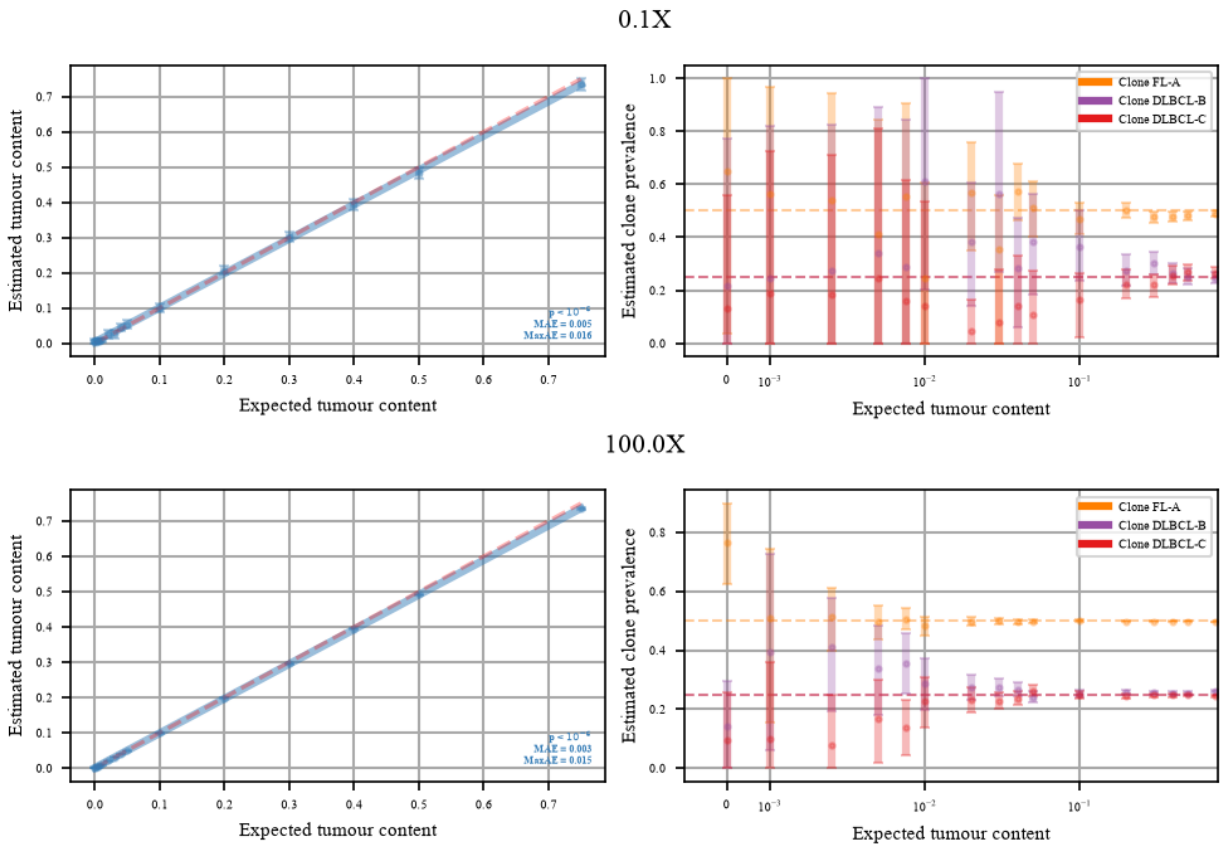

Figure 13: Tumour content (left) and clone prevalence (right) estimation with 95% HDI intervals on data generated with the infinite pool model using scWGS data from FL and DLBCL cancers and a matched normal sample. Coverages varied for 8 values between $0.1X$ and $100X$ and tumour content was varied for 16 values between 0% and 75%. True clone prevalence are indicated by the dashed line where Clone A, Clone B, and Clone C have true values of 0.5, 0.25, and 0.25.

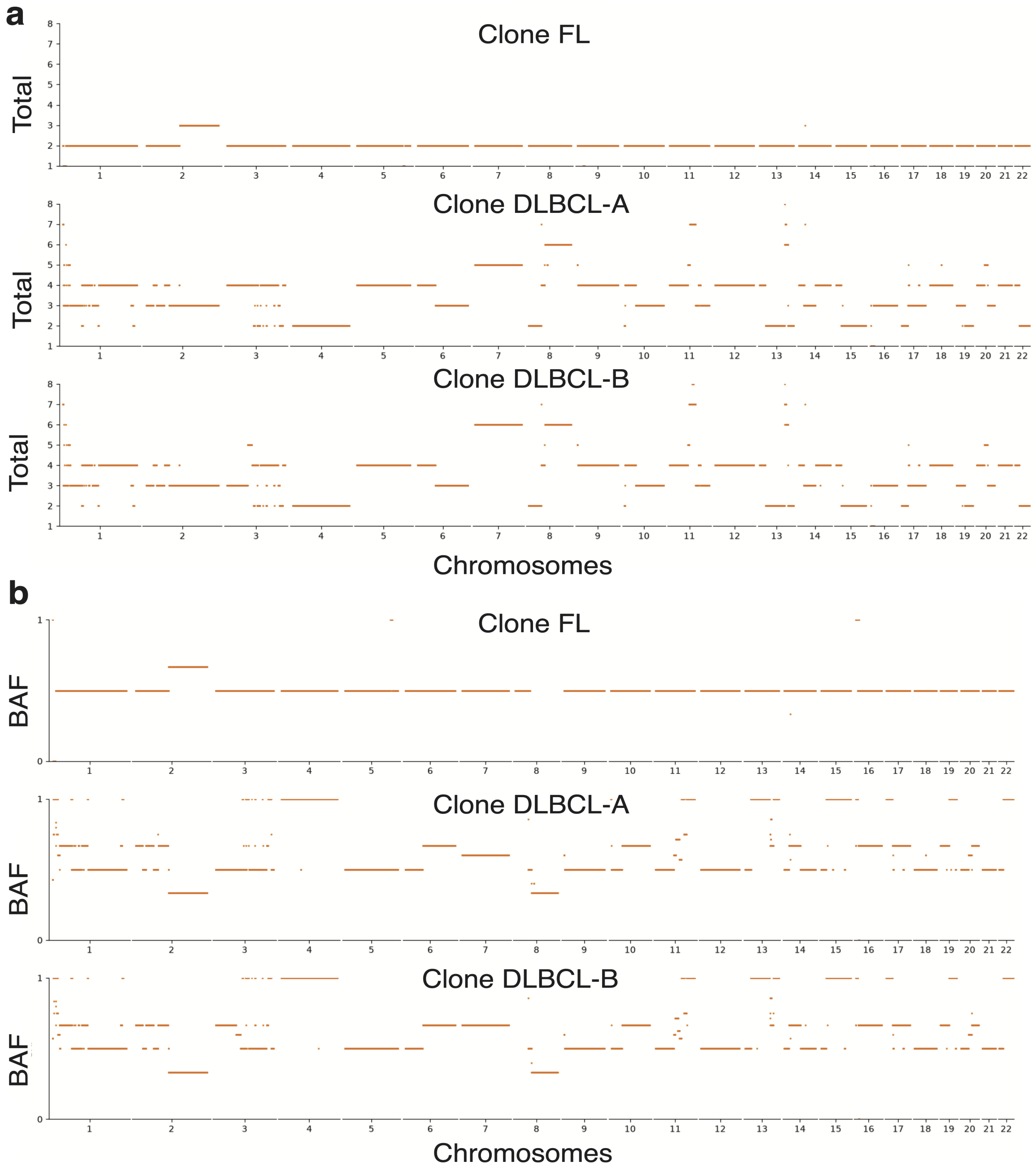

Figure 14: Clone total copy-number (a) and BAF (b) profiles for the FL and DLBCL clones obtained from scWGS.

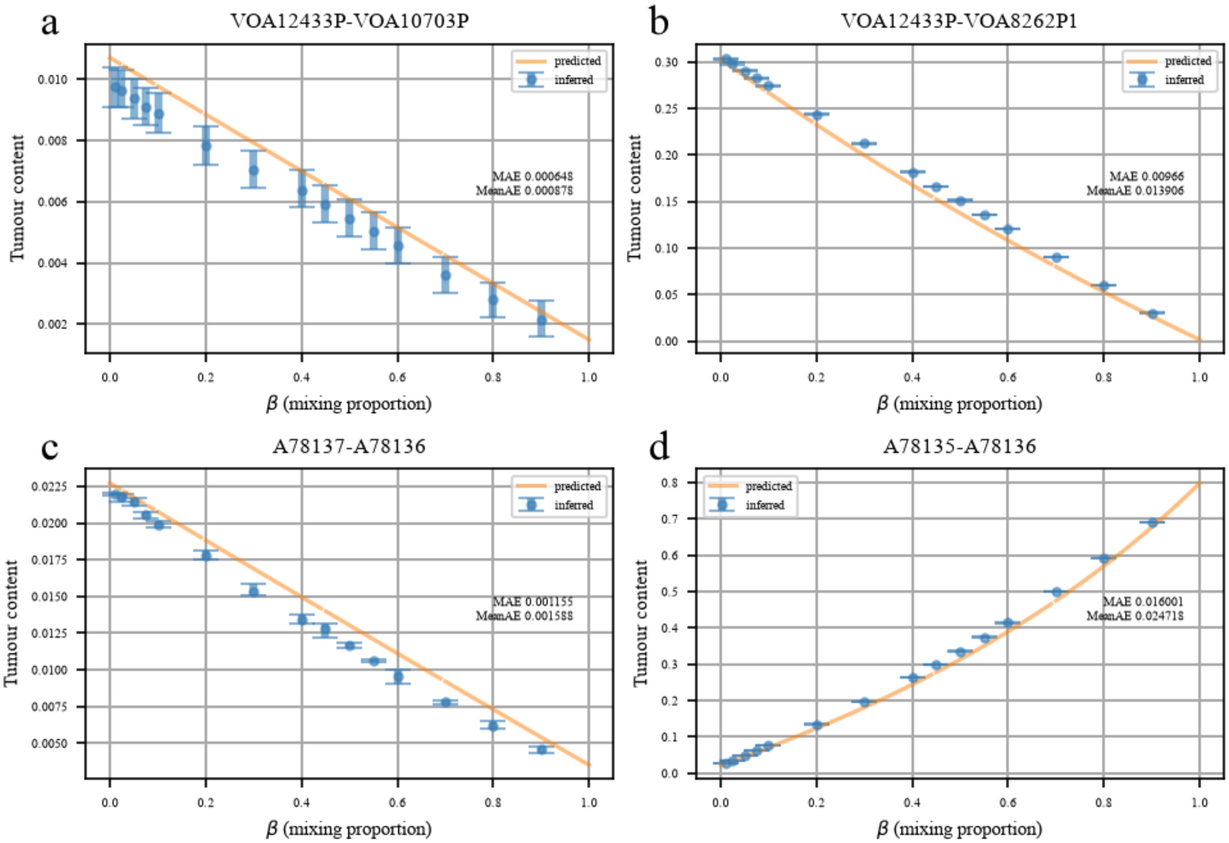

Figure 15: Inferred mean tumour fraction and 95% HDI interval from cfClone (blue) and predicted tumour fraction by mathematical model (orange) for in-silico mixtures generated at $30X$ by mixing patient derived cfDNA containing zero estimated tumor content with cfDNA estimated to contain tumor content (**a**, **b**) and tissue (**c**, **d**) samples with variable estimated tumor content.

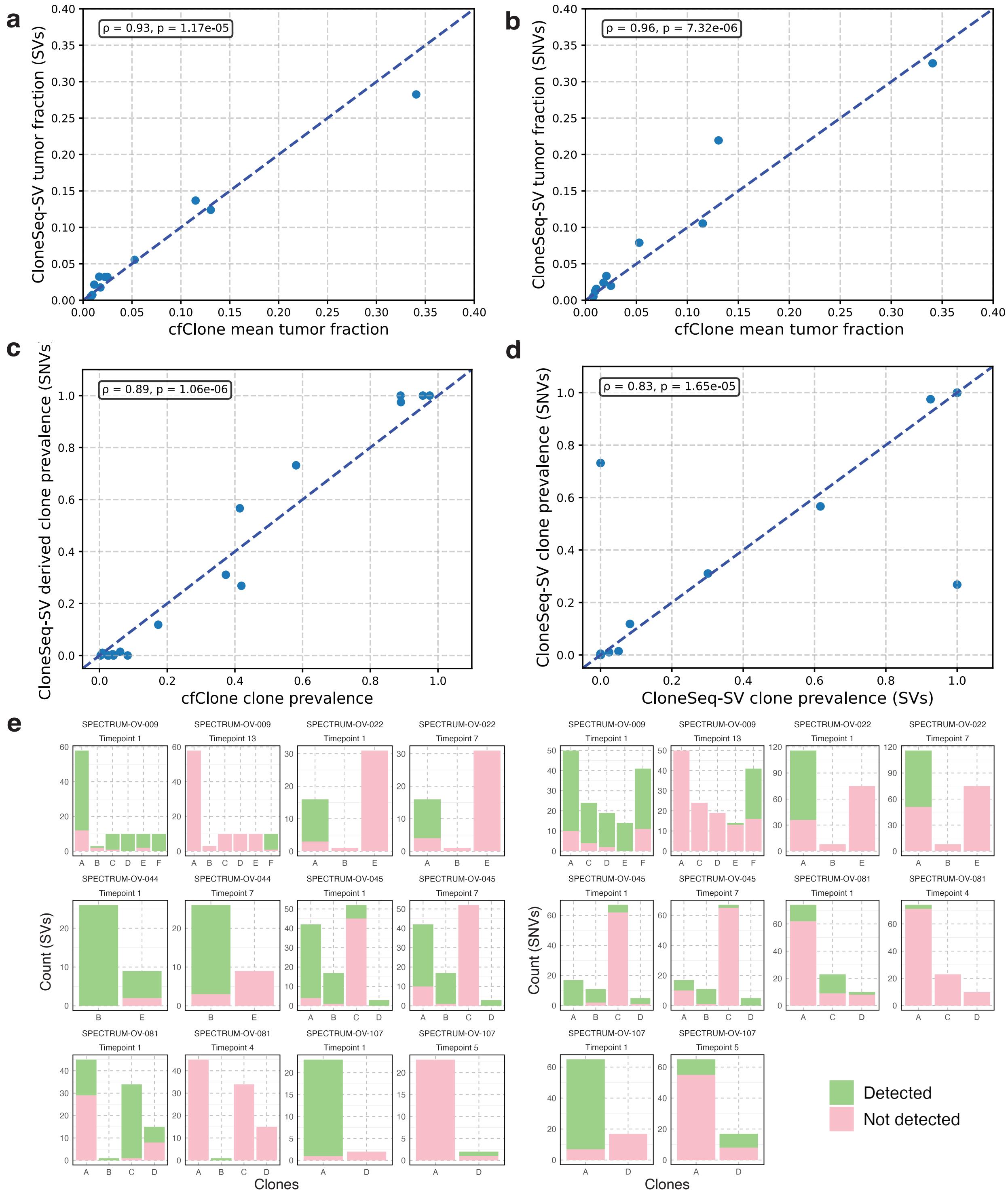

Figure 16: Validation on the MSK-SPECTRUM dataset **a**, The mean tumor fraction per sample estimated by cfClone versus the tumor fraction estimated by CloneSeq-SV using SVs (n = 6 patients, 12 samples). **b**, The mean tumor fraction per sample estimated by cfClone versus the tumor fraction estimated by CloneSeq-SV using SNVs (n = 5 patients, 10 samples). **c**, The mean clonal prevalence estimated by cfClone versus the clonal prevalence estimated by CloneSeq-SV using SVs (n = 3 patients, 6 samples). **d**, The clonal prevalence estimated by CloneSeq-SV using SVs versus the clonal prevalence estimated by CloneSeq-SV using SNVs (n = 3 patients, 6 samples). **e**, The number of SVs and SNVs detected at each timepoint by CloneSeq-SV.

### List of Tables

1 Notation used in the cfClone model. . . . . 31
2 Mean and max absolute error of tumour fraction estimates for cfClone and ichorCNA for three different cancer types FL, DLBCL, and HGSOC. Metrics are computed using estimates on data generated using 16 values of tumour content between 0. and 0.75 for coverage $100X$. . . 32

Table 1: Notation used in the cfClone model.

| **Symbol** | **Range** | **Description** |
|---|---|---|
| $t$ | $[T]$ | Genomic bin index |
| $k$ | $[K+1]$ | If $k \in [K]$, the index corresponds to a clone; if $k = K+1$ the index corresponds to the normal cell population |
| $L_t$ | $(0, \infty)$ | RDR for genomic bin $t$ |
| $D_t$ | $\{0, 1, 2, \dots\}$ | Total number of reads overlapping heterozygous SNPs in genomic bin $t$ |
| $B_t$ | $\{0, 1, 2, \dots, D_t\}$ | Number of reads mapping to haplotype B in bin $t$ |
| $\mu_t$ | $(0, \infty)$ | Predicted RDR location parameter for bin $t$ |
| $p_t$ | $[0, 1]$ | Predicted haplotype B frequency for genomic bin $t$ |
| $\rho$ | $[0, 1]^{K+1}$ | Probability vector of size $K+1$ denoting normal and clonal prevalences, $\rho = (\rho_1, \rho_2, \dots, \rho_{K+1})$, $\sum_k \rho_k = 1$ |
| $\sigma$ | $(0, \infty)$ | Scale parameter of the RDR likelihood model |
| $\alpha$ | $(0, \infty)$ | Shift parameter used to compute the RDR predicted location; deviation from $\alpha = 1$ provides robustness to non-uniform coverage |
| $\pi_{\text{rdr}}$ | $[0, 1]$ | Mixture probability for the outlier distribution in the RDR likelihood model |
| $\pi_{\text{baf}}$ | $[0, 1]$ | Mixture probability for the outlier distribution in the BAF likelihood model |
| $\gamma$ | $(0, 1)$ | "Divergence from binomiality" parameter for the BAF likelihood model |
| $C_{tk}$ | $\{0, 1, 2, \dots\}$ | Total copy number of clone $k$ in genomic bin $t$ |
| $C_{tk}^b$ | $\{0, 1, 2, \dots, C_{tk}\}$ | Haplotype B copy number for clone $k$ in genomic bin $t$ |
| $C_{tk}^a$ | $\{0, 1, 2, \dots, C_{tk}\}$ | Haplotype A copy number for clone $k$ in genomic bin $t$ |

| | Mean absolute error | | Max absolute error | |
|---|---|---|---|---|
| | cfClone | ichorCNA | cfClone | ichorCNA |
| **FL** | 0.2% | 0.3% | 0.8% | 3% |
| **DLBCL** | 0.2% | 2.4% | 0.9% | 22% |
| **HGSOC** | 0.1% | 1.7% | 0.7% | 5% |

Table 2: Mean and max absolute error of tumour fraction estimates for cfClone and ichorCNA for three different cancer types FL, DLBCL, and HGSOC. Metrics are computed using estimates on data generated using 16 values of tumour content between 0. and 0.75 for coverage $100X$.
